# Membrane-Promoted J-Aggregation of BODIPY Dimers Enables Spontaneous Blinking in Green and Red Channels for Live-Cell Nanoscopy

**DOI:** 10.64898/2026.01.19.700264

**Authors:** Sonia Pfister, Sophie Walter, Pascal Didier, Mayeul Collot

**Author notes:** **Corresponding Author** Dr. Mayeul Collot. Chemistry of Photoresponsive Systems, Laboratoire de Chémo-Biologie Synthétique et Thérapeutique (CBST) UMR 7199, CNRS, Université de Strasbourg, F-67400 Illkirch, France.

## Abstract

BODIPY dyes are widely used in bioimaging, yet their aggregation behavior within the plasma membrane (PM) remains poorly exploited for single-molecule localization microscopy (SMLM). Here, we design a series of green-emitting PM-targeted BODIPY dimers engineered to undergo spontaneous and transient *H*- and *J*-aggregation. By tuning the linker length between the fluorophores, we identify dimers that form intramolecular *H*-aggregates in polar media and emissive *J*-aggregates (λ_em_ ≈ 535 nm) through membrane-driven intermolecular interactions. In lipid bilayers, all dimers aggregate above a probe/lipid ratio of 1/100, exclusively generating *J*-aggregates. In live-cell SMLM, the monomeric MB-488 provides high event numbers via diffusion-driven emission, whereas dimers exhibit stable blinking and yield brighter red-shifted *J*-aggregate events with improved localization precision. Red-channel events localize to specific PM regions, suggesting preferential *J*-aggregation within distinct membrane microdomains. HaloTag constructs targeted to cell-surface PDGFR further confirm that intramolecular *J*-aggregation is possible but strongly amplified by the membrane through intermolecular collisions. These results demonstrate that green BODIPYs and their *J*-aggregates enable robust live SMLM in green and red channels, and reveal the PM as a privileged environment for emissive *J*-aggregation.

## Introduction

Too often reduced to a simple phospholipid bilayer, the plasma membrane (PM) is in fact a highly organized and dynamic structure in which are anchored various receptors and channels that govern important biological events and the communication of the cell with its environment. Therefore, high-fidelity PM bioimaging has become a major challenge. Owing to its excellent spatial and temporal resolution and its compatibility with live cells, fluorescence microscopy has emerged as a central technique and is now widely accessible. Consequently, chemists have focused their efforts on developing fluorescent probes selective for the PM and capable of sensing variations in polarity, viscosity, temperature, or tension.^1^ Yet, simple and efficient PM stains remain invaluable for cell segmentation and for visualizing thin PM protrusions such as filopodia^2^ or tunneling nanotubes (TNTs).^3 4 5^

In this context we developed MemBright®, a family of probes that rapidly and efficiently stain live and fixed cells^4,6^ which has since gained widespread adoption. The advent of super resolution fluorescence techniques such as Stimulated Emission Depletion microscopy (STED),^7 8 9^ and Single Molecule Localization Microscopy (SMLM),^10 11^ has intensified the need for PM probes tailored for these advanced imaging modalities.^12^ Fluorescent probes enabling SMLM imaging of the PM in fixed cells have been reported.^12^ Within the MemBright family, MBCy3.5, a red-emitting probe, proved particularly effective for STORM imaging, notably of fragile neuronal cells.^6, 13^ However, live-cell-compatible probes remain scarce as SMLM demands the rapid collection of numerous spatially distributed localization events before significant sample motion occurs.

One strategy to meet these constraints is to capture the signal of a fluorescent molecule diffusing within the lipid bilayer,^14 15 16^ an approach that has enabled quantitative mapping of membrane diffusivity and polarity in live cells.^10^Another powerful strategy is PAINT (Point Accumulation for Imaging in Nanoscale Topography), which is particularly well suited for live-cell PM super-resolution imaging, as it relies on transient and fluorogenic probe–membrane interactions.^11, 17, 18^ Conversely, PM-anchored photoswitchable probes that undergo light-induced changes in their photophysical properties can also support live-cell SMLM.^19^ While spirolactamization remains the most widespread spontaneously blinking mechanism,^20–24^ we recently showed that, when confined to the PM, the local hydrophobicity dramatically alters the effective pKa of these probes and thus their switching behavior.^25^ In a distinct approach, we introduced the concept of directed photooxidation enabling the design of photoconvertible,^26^ and photoactivatable^27^ BODIPYs suitable for efficient live-cell SMLM notably at the PM.^26^ Also built on photooxidation, we recently introduced the concept of Self-Triggered Photooxidation Cascade (STPC) in which a leuco-rhodamine serves as a photoactivatable PM probe which enabled super-resolution microscopy of live cells’ PM, allowing super-resolution imaging of the PM and time laps imaging of thin PM processes.^5^

In principle, molecular switching may also arise from the reversible formation of non-emissive *H*-aggregates. This concept has been exploited in conventional laser scanning confocal microscopy to design fluorogenic oligomeric,^28^ and more commonly, dimeric probes for wash-free, high signal to noise imaging of various membrane receptors using green BODIPY,^29^ squaraine dyes,^30 31^ or cyanines,^32, 33^ as well as for RNA-aptamer using rhodamines.^34^

Recently, a PAINT-inspired strategy employing Cy3 and Cy5 dimers with low PM affinity was introduced to achieve reversible PM binding.^18^ This approach enabled 1) a substantial reduction of background signal due to aggregation-caused quenching (ACQ) in the aqueous extracellular environment and 2) enhanced single-molecule brightness resulting from the simultaneous activation of two fluorophores instead of one. Despite these efficient approaches, the formation of aggregates within the PM has never been explored as a means to generate spontaneous fluorescence blinking in live SMLM. Unlike *H*-aggregates, *J*-aggregates are reported as red-shifted emitting species and could thus provide a reversible spectral shift. Despite these attractive features, *J*-aggregation remains largely underexplored in bioimaging. Indeed, although *J*-aggregation of BODIPYs is widely used to develop nanomaterials with red-shifted emission for *in vivo* imaging,^35, 36, 37, 38, 39, 40^ transient molecular *J*-aggregation has virtually never been investigated as a switching mechanism for SMLM applications. Only recently, Adhikari *et al*. showed that emissive BODIPY fatty-acid analogues could transiently form red-shifted intermolecular aggregates assigned to *J*-aggregation, and elegantly used this approach for multicolor SMLM experiments. ^41^

In this work, we investigated the ability of BODIPY dimers to undergo transient *H-* and *J*-aggregation once embedded in the PM, thereby enabling fluorescence switching in a green and red channel, respectively, for live-cell super resolution cell imaging (Figure 1).

**Figure 1.**
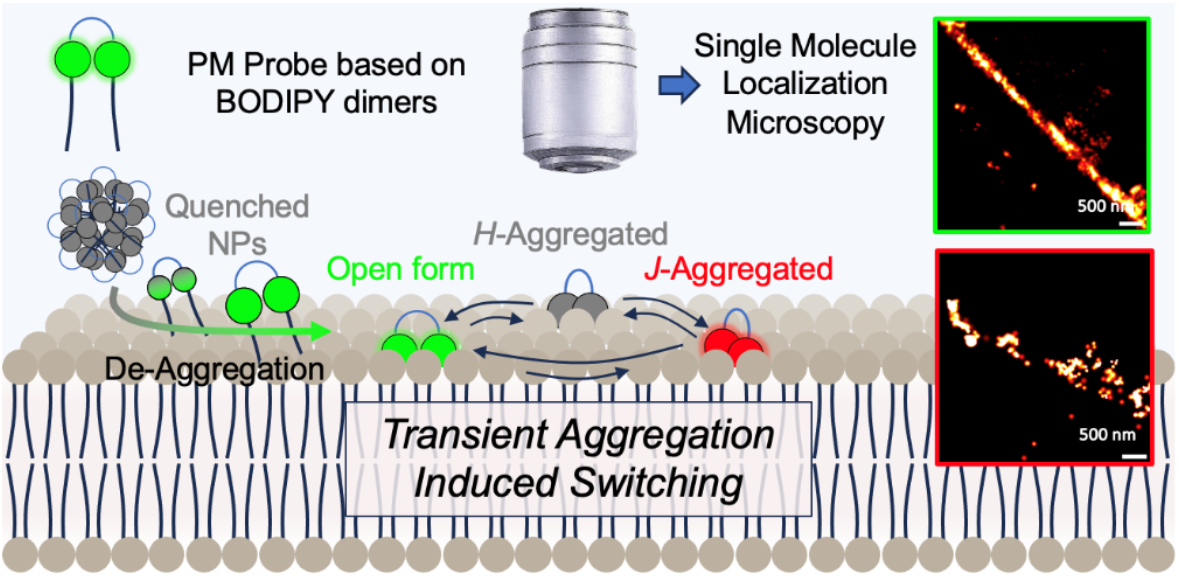
Plasma membrane (PM) probes based on BODIPY dimers were synthesized and evaluated for their ability to form transient *H*- and *J*-aggregates. The dynamic equilibrium between the emissive open form and the non-emissive *H*-aggregated state, as well as the red-shifted *J*-aggregated state, induces spontaneous fluorescence switching at the PM, enabling live-cell Single Molecule Localization Microscopy imaging.

## Results and discussion

### Design

Simple and “conventional” green-emitting BODIPY were chosen for their relative photostability, the robustness of their photophysical properties across diverse environments,^42^ and their ability to form both *H*-,^4, 43^ and *J*-aggregates.^41, 44–46^ To evaluate the spontaneous blinking behavior of BODIPY dimers in the PM, several probes were designed (Figure 2). All of them possess two amphiphilic anchors that have previously demonstrated high efficiency as PM-targeting moieties.^4, 5, 6, 9, 25, 26, 47^ To access different aggregation patterns (*H*-vs. *J*-aggregation), spacers of increasing length were introduced. First, the BODIPY Dimer–Lysine (BD-Lys) was designed to provide a minimal separation between the two BODIPYs. Lysine has previously been shown to efficiently promote *H*-aggregation in related dye architectures.^28, 29, 30, 31, 34^ To provide additional conformational freedom to the dimers, PEG linkers of increasing lengths (from PEG4 to PEG12) were used to separate the two BODIPYs, each bearing an amphiphilic PM targeting unit. Importantly, MemBright-488 (MB-488), an efficient PM that we developed,^4^ was used as a monomeric control to assess the specific effects arising from PM-embedded BODIPY dimers.

**Figure 2.**
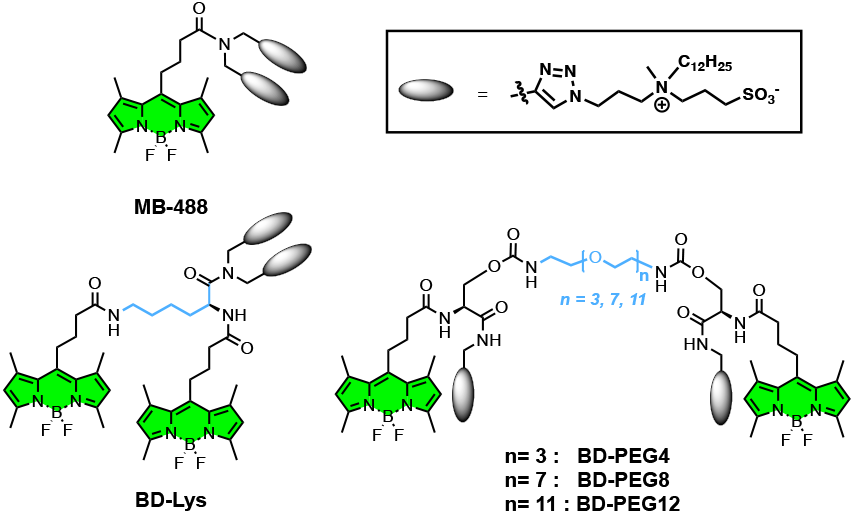
Chemical structures of the monomeric PM probe Mem-Bright-488 (MB-488) and the four BODIPY dimers (BD-Lys, BD-PEG4, BD-PEG8, BD-PEG12) designed to modulate spacing and aggregation propensity.

### Synthesis

The synthetic strategy for the dimeric PM probes involved an initial preparation of the BODIPY dimer precursors (p-BD), followed by their conversion into their corresponding plasma membrane probes (BD) via CuAAC (scheme 1). BD-Lys was synthesized in three steps. Di-Boc-protected lysine was first di-propargylated, then the Boc groups were removed under acidic conditions to afford intermediate 1. This compound was then coupled with BDP-COOH to yield the first BODIPY dimer precursor, p-BD-Lys. PEG-separated dimers were obtained from the propargylation of *N*-Boc-L-serine, followed by Boc deprotection. The resulting amine 2 was coupled with BDP-COOH to furnish alcohol 3. The hydroxyl group of 3 was then converted into an activated carbonate, giving rise to key intermediate 4. PEG chains of various lengths (PEG4, PEG8, PEG12) were subsequently coupled to intermediate 4 to afford the corresponding precursors p-BD-PEG4, p-BD-PEG8, and p-BD-PEG12. In the final step, all BODIPY dimer precursors were clicked to the clickable amphiphilic zwitterion (CAZ) targeting moiety, yielding the final dimeric PM probes (scheme 1).

**Scheme 1.**
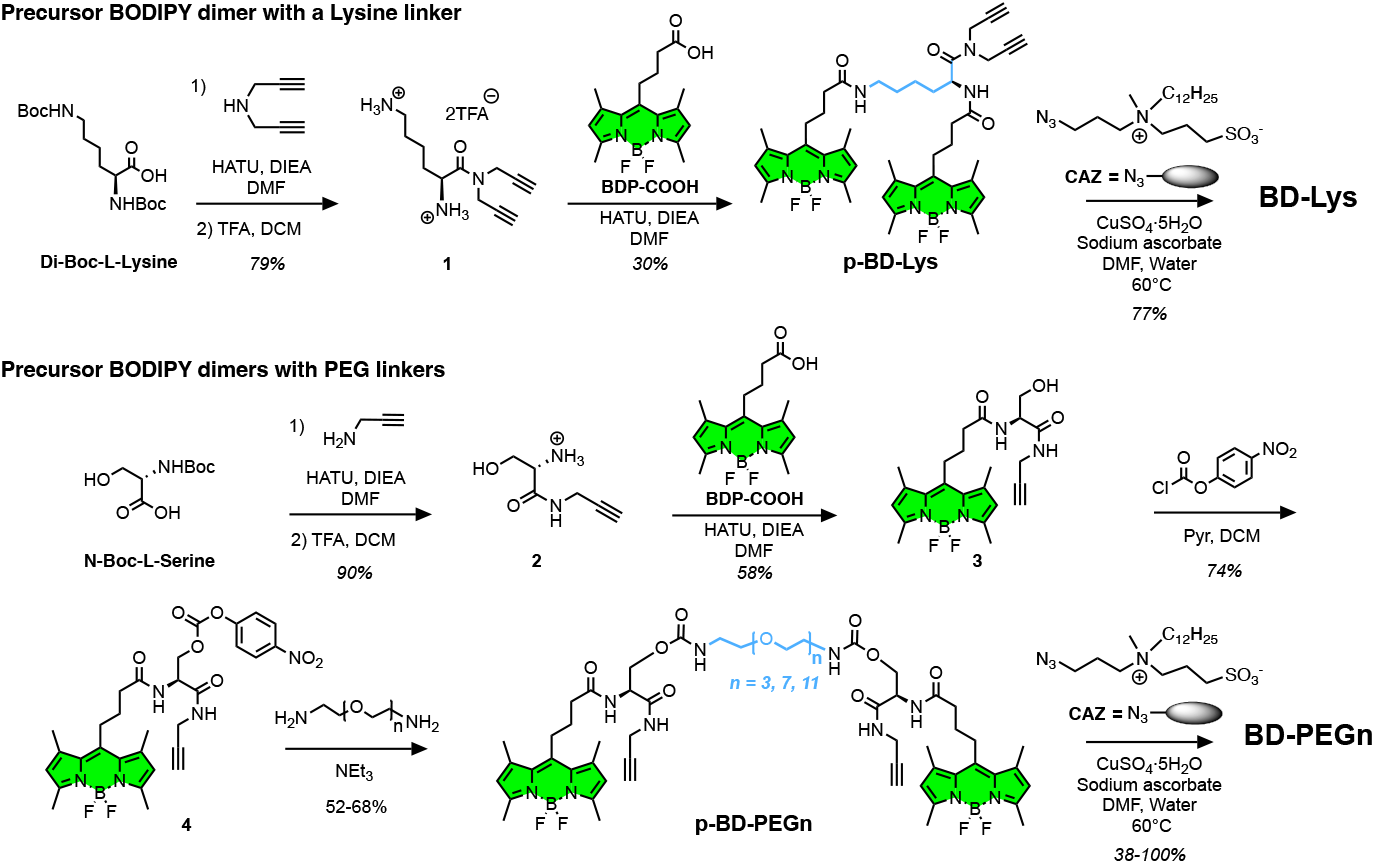
Synthesis of the precursors BODIPY dimers (p-BD) and their corresponding plasma membrane probes (BD).

### Photophysical properties

The photophysical properties of the BDs were evaluated in methanol, where the probes are fully soluble, in phosphate buffer (pH 7.4) to mimic the extracellular environment, and finally in DOPC large unilamellar vesicles (LUVs in PBS) as plasma membrane (PM) models. The results are reported in Table 1 and Figure 3. As expected, the dimers displayed typical maximum absorption and emission wavelengths centered around 500 and 506 nm, respectively, with low Stokes shifts (5 to 9 nm).

**Table 1.** Photophysical properties of the PM dimeric probes BDs in different environments.

| Solvent | Probe | $\lambda_{abs\ max}$<br>(nm) | $\lambda_{em\ max}$<br>(nm) | $\Phi_{FI}$ |
| --- | --- | --- | --- | --- |
| MeOH | BD-Lys | 497 | 506 | 0.58 |
|  | BD-PEG4 |  |  | 0.60 |
|  | BD-PEG8 |  |  | 0.62 |
|  | BD-PEG12 |  |  | 0.50 |
| PBS<br>(pH = 7.4) | BD-Lys | 501 | 506 | 0.02 |
|  | BD-PEG4 |  |  | 0.03 |
|  | BD-PEG8 |  |  | 0.05 |
|  | BD-PEG12 |  |  | 0.04 |
| DOPC LUV<br>in PBS<br>(pH = 7.4) | BD-Lys | 499 | 507 | 0.94 |
|  | BD-PEG4 |  |  | 0.99 |
|  | BD-PEG8 |  |  | 0.86 |
|  | BD-PEG12 |  |  | 0.86 |

**Figure 3.**
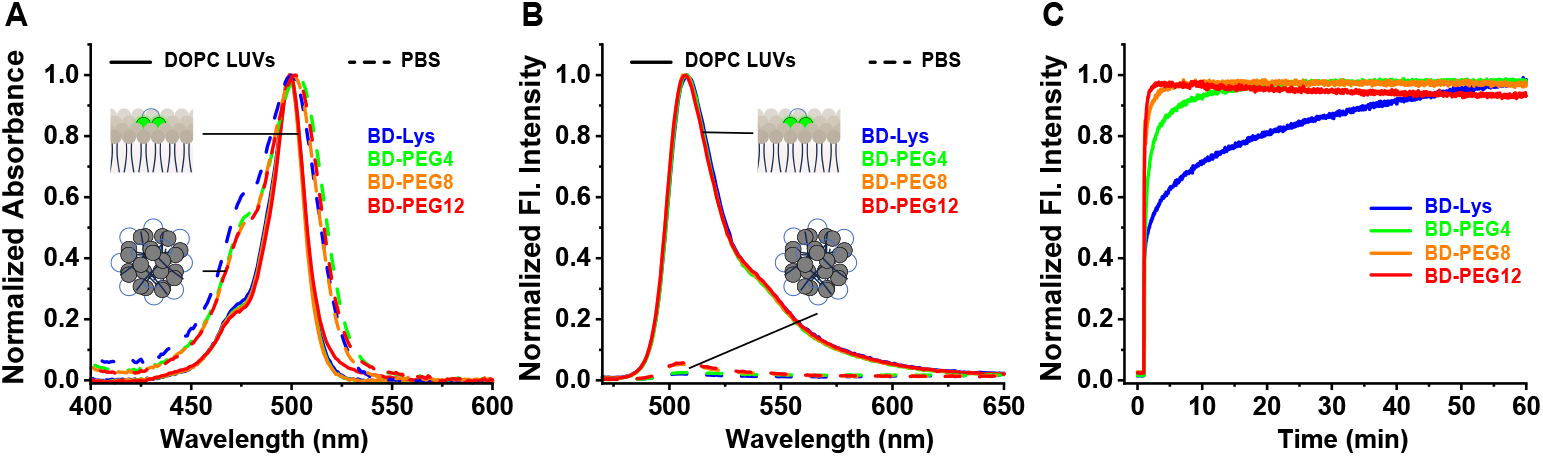
Fluorogenic properties of the dimer-based PM probes. (A) Normalized absorption spectra of the BD in PBS and embedded in DOPC LUVs. (B) Normalized emission spectra in the same conditions; the spectra in PBS were normalized to the LUVs maxima to highlight the fluorescence enhancement. (C) Kinetics of fluorescence enhancement upon addition of probes (1 µM) to DOPC LUVs (200 µM). λ_exc_ = 460 nm.

In methanol,the probes are fully solubilized and display fluorescence quantum yields ranging from 0.50 to 0.62. Conversely, in aqueous media, the dimers exhibit low quantum yields (0.02– 0.05) due to aggregation-caused quenching (ACQ), driven by their amphiphilic nature. ACQ was confirmed by the broadening of the absorption spectra, indicative of aggregation (Figure 3A). Upon incorporation into phospholipid bilayers, the absorption spectra recovered their regular shape (Figure 3A) and were accompanied by high quantum yield values, suggesting that in all cases the probes are de-aggregated and the fluorophores are well separated once inserted into the plasma membrane model. The pronounced fluorescence enhancement observed between aqueous media and the PM model (up to 47-fold) makes these dimers efficient fluorogenic PM probes (Figure 3B).

Interestingly, dimers with long separations, BD-PEG8 and BD-PEG12, display lower quantum yields (0.86) than those with short linkers, namely BD-Lys and BD-PEG4 (0.94 and 0.99, respectively). This observation might be attributed to the increased polarity and water-solubilizing effect of long PEG chains, which can transiently expose the dimer to the aqueous phase, thereby decreasing its brightness.

This difference of polarity between the dimers was also reflected in their binding kinetics on LUVs (Figure 3C). Whereas the relatively hydrophobic BD-Lys binds in a slow manner, PEG-separated dimers bind rapidly to LUVs according to their increasing polarity (*i*.*e*., PEG length) with binding kinetics following BD-PEG12 > BD-PEG8 > BD-PEG4. Indeed, we already showed that PM probes, due to their amphiphilic nature form large aggregates in aqueous media and that the more polar the PM probes, the faster they de-aggregate and insert into lipid membranes.^4, 6^ Overall, the BD PM probes possess fluorogenic properties which are similar to their monomer cognate MB-488.^4^

### Investigation on aggregation patterns

To study the effect of linker length, which separates the two fluorophores, on aggregation behavior, the absorption spectra of the dimers were recorded in methanol with increasing proportions of water in order to progressively induce aggregation. Indeed, absorption spectra provide valuable insight into the aggregation state of fluorophores. While spectral broadening mainly reflects intermolecular aggregation, the appearance of a hypsochromically or bathochromically shifted band corresponds to *H*- and *J*-aggregation, respectively. For this purpose, precursor BODIPY dimers (p-BDs) were used instead of the final amphiphilic dimers, as the latter rapidly undergo intermolecular aggregation due to their amphiphilic character.

Interestingly, the p-BDs exhibited markedly different changes in their absorption spectra, and thus in their aggregation patterns, upon increasing water content (Figure 4). Whereas p-BD-Lys showed signs of intramolecular *H*-aggregation as water content increased, evidenced by narrow spectra (FWHM = 21 nm) and an enhanced blue-shifted shoulder at 475 nm, its spectra began to broaden significantly (up to FWHM = 58 nm) from 80% water, indicating the onset of intermolecular aggregation (Figure 4A). Conversely, the more polar p-BD-PEG12, in which the fluorophores are more widely separated, progressively formed intramolecular *H*-aggregated dimers even at high water content (Figure 4D). This behavior was attributed to the strong solubilizing effect of the long PEG12 chain, which prevents intermolecular aggregation and instead promotes intramolecular dimerization mediated *H*-aggregation.^30^

**Figure 4.**
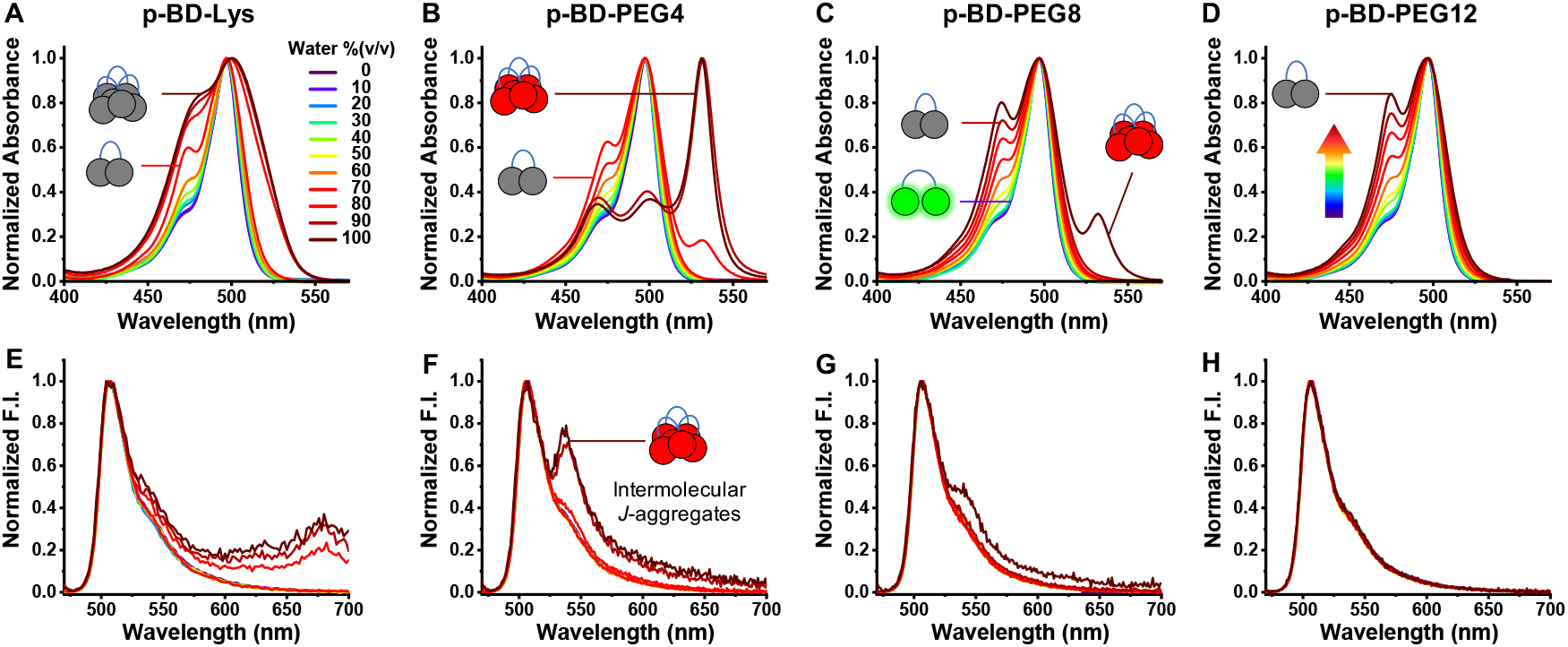
Aggregation of precursor p-BDs in MeOH/H_2_O mixtures. (A–D) Normalized absorption spectra showing emergence of *H*-aggregates (blue-shifted shoulder) and *J*-aggregates (narrow and red-shifted band). (E–H) Corresponding emission spectra. p-BDs concentration was 1 µM. Green, grey, and red dimers represent respectively: non-aggregated open form, intra- and inter-molecular *H*-aggregates, and intermolecular *J*-aggregates.

PEG-separated dimers of intermediate linker length, namely p-BD-PEG4 and p-BD-PEG8, initially displayed the same intra-molecular *H*-aggregation trend as the other p-BDs. However, at high water contents (80% and 100% for p-BD-PEG4 and p-BD-PEG8, respectively), their absorption spectra exhibited a narrow (FWHM = 16 nm) red-shifted band with λ_abs max_ = 532 nm, characteristic of *J*-aggregation.

To determine whereas these *J*-aggregates were the result of intramolecular or intermolecular aggregation, absorption spectra in high water content was performed with increasing concentration (Figure S1). The results showed that the red-shifted *J*-band decreased at low concentrations for p-BD-PEG4, whereas it disappeared upon dilution for p-BD-PEG8, suggesting that inter-molecular aggregation mainly contributes to *J*-aggregate formation for these dimers.

At this stage, the same experiments were performed with the BODIPY monomer p-MB488, the precursor of MB-488 (FigureS2). Its absorption spectra indicated only a weak tendency to form *H*-aggregates, even in 100% water and at high concentration up to 6 µM.

In parallel fluorescence emission spectra were recorded. As expected, all BDs exhibited a decrease in fluorescence intensity as water content increased (Figure S3). In agreement with the absorption data, the emission spectra (Figure 4E–H) revealed that whereas p-BD-PEG12 retained its usual spectral signature, indicative of classical ACQ, p-BD-PEG4 was the most prone to form emissive *J*-aggregates, as shown by the appearance of a narrow red-shifted emission band (λ_em max_ = 535 nm) at 90% water (v/v). The characterization of p-BD-PEG4 *J*-aggregates proved challenging due to their small Stokes shift (4 nm, figure S4), their low spectral shift compared to the non aggregated form and their weak emission, while a significant fraction of non-aggregated p-BD-PEG4 remained highly emissive. We ultimately succeeded in reconstructing a reliable emission spectrum (see Materials and Methods and Figure S4). p-BD-PEG4 *J*-aggregates emit at 535 nm as a narrow band (FWHM = 16 nm). Although the excitation spectrum of p-BD-PEG4 *J*-aggregates overlapped with that of the remaining bright green non-aggregated dimers (band at 497 nm), it clearly displayed the red-shifted band at 531 nm, matching the band observed in absorption. This demonstrates that the red-shifted absorption band is responsible for the 535-nm emission, thus confirming the formation of emissive *J*-aggregates.

Notably, in the literature,^41, 44^ *J*-aggregates of BODIPY dyes are often observed at even more strongly red-shifted wavelengths, highlighting the unusually compact packing and specific geometry of the present *J*-aggregates

### Aggregation in membrane models

Having shown that precursor BODIPY dimers (p-BDs) readily form *H*- and *J*-aggregates in highly polar media, we next investigated whether their corresponding plasma membrane probes (BDs) could also form aggregates once embedded within a lipid bilayer. To this end, BDs were incorporated into DOPC LUVs at increasing probe/lipid ratios in order to promote aggregation by raising their local concentration (Figure 5). Except for BD-PEG12, the absorption spectra revealed that increasing the local probe concentration induced a red shift accompanied by spectral broadening. The fluorescence spectra exhibited similar behavior, with the appearance of a red-shifted broadened band, particularly pronounced for BD-PEG4 and BD-PEG8. Notably, we found that for all probes, the linear increase of fluorescence intensity abruptly broke at 2 μM (Figure 5E–H, insets), indicating that aggregation begins to occur in DOPC LUVs above a probe/lipid ratio of 1/100. Interestingly, although the break in linearity clearly signaled aggregation, the absorption spectra of the BDs in lipid bilayers showed no evidence of *H*-aggregation (Figure 5E–H). These observations suggest that the observed decrease in fluorescence intensity is primarily due to the formation of weakly emissive *J*-aggregates, rather than *H*-aggregates. Under the same conditions, the monomeric PM probe MB-488 did not show significant changes in its absorption spectra. However, at a probe/lipid ratio of 1/20, its emission spectrum began to red-shift (Figure S5), suggesting that MB-488 can also form *J*-aggregates within lipid bilayers but at higher local concentration.

**Figure 5.**
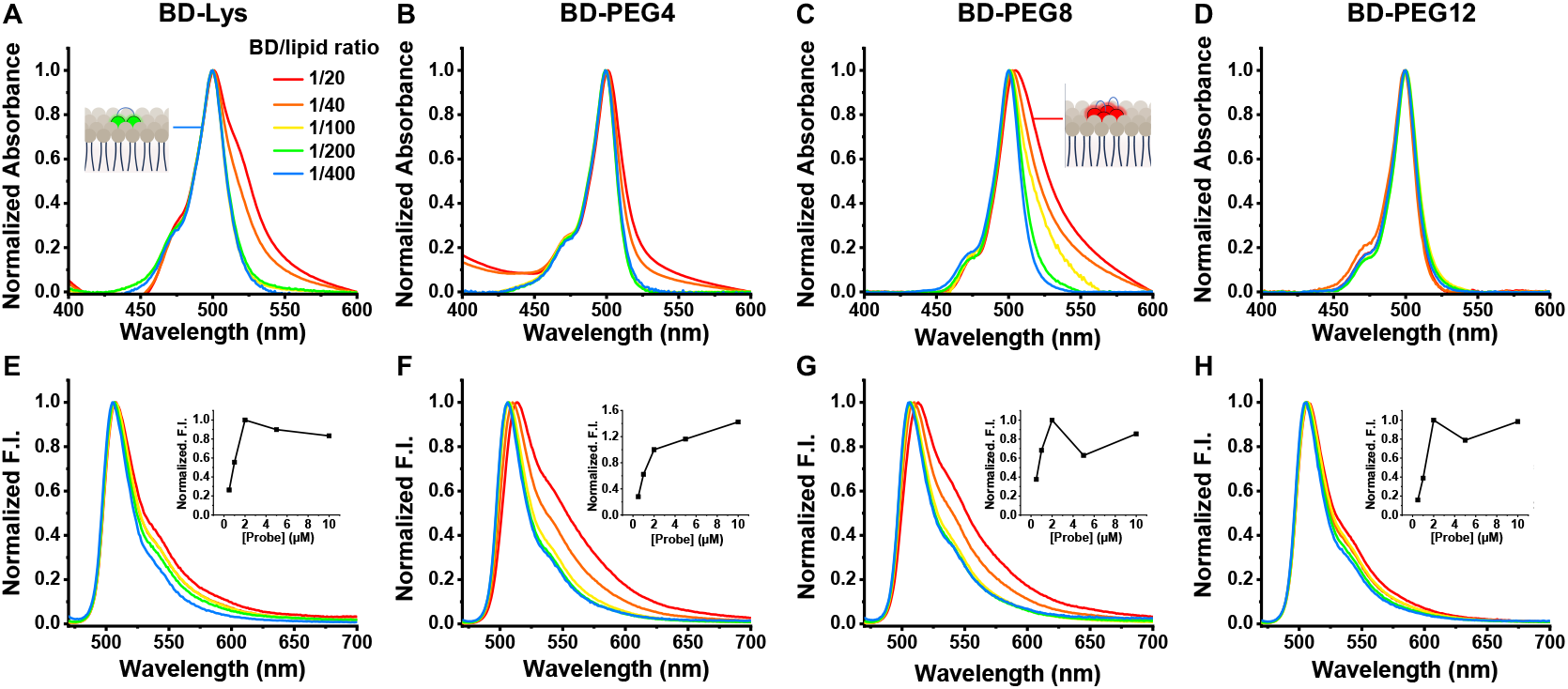
Aggregation of membrane-embedded BDs in DOPC LUVs (in PBS pH 7.4). Normalized absorption (A-D) and emission (E-H) spectra of BDs PM probes once inserted in DOPC LUVs at various Probe/Lipid ratio to increase their local membrane concentration. The concentration of lipids was fixed at 200 µM and the concentration of probe was increased from 0.5 to 10 µM, providing the probes/lipid ratio indicated in A.

Taken together, these results show that, unlike the monomer, BODIPY dimers can form both *H*- and *J*-aggregates. As expected, and as demonstrated in our previous work, dimerized BODIPYs efficiently form intramolecular *H*-aggregates in polar environments. Here, we show that depending on the nature and length of the linker between the dyes, green BODIPY dimers can also form emissive, red-shifted *J*-aggregates. Notably, in our model, PEG4 and PEG8 appear to be the most suitable linkers, providing an optimal balance of rigidity and flexibility to enable fluorophore orientations conducive to *J*-aggregation.

In lipid bilayers, the aggregation pattern differs: no *H*-aggregation is observed in absorption, while the red shift of both absorption and emission spectra clearly indicates *J*-aggregate formation, which is in line with a previous work.^48^ The concentration-dependent experiments further revealed that *J*-aggregates arise not from intramolecular dimer folding, but rather from intermolecular interactions. Indeed, dimerized BODIPYs behave differently from monomeric MB-488, forming *J*-aggregates at lower concentrations and to a much greater extent. These results therefore indicate that dimerization strongly promotes *J*-aggregation through intermolecular interactions.

Finally, although ensemble spectroscopic data indicate that dimers preferentially form *J*-aggregates, the rare and transient formation of *J*-aggregates by monomeric species may still be sufficient, at the single-molecule scale, to enable SMLM imaging. Based on the work of Adhikari *et al*. on transient red-shifted aggregates formed by single BODIPY molecules in membranes,^41^ we propose that BODIPY dimers may transiently alternate between an open form and a *J*-aggregated form within the membrane, thereby producing efficient blinking in the green and red microscopy channels.

## CELLULAR STUDIES

Prior to SMLM experiments, the cytotoxicity of the probes was evaluated using MTT cell viability assays. The results showed no significant toxicity (Figure S6). Next, the ability of the BDs to stain the PM in a selective manner was assessed. Live HeLa cells were stained with BDs along with MemBright-Cy5.5 as a counterstain (Figure S7). The cells were imaged without washing after 10 min and showed: (1) selective PM staining, as demonstrated by the high Pearson colocalization coefficients, and (2) intense PM fluorescence in the green channel, with a high signal-to-noise ratio due to the strong fluorogenic effect of the probes.

### Live single molecule localization microscopy

To evaluate the blinking ability in the PM of live cells, we first performed SMLM experiment at increasing probe concentrations (from 0 to 250 nM) and measured the average number of events per PM surface in the green channel (Figure 6A). A significant increase in the number of events was observed at 50 nM; however, higher concentrations did not improve event counts and even decreased performance. Surprisingly, at this optimal concentration of 50 nM, no major performance differences were observed between the different PEG-containing dimers.

**Figure 6.**
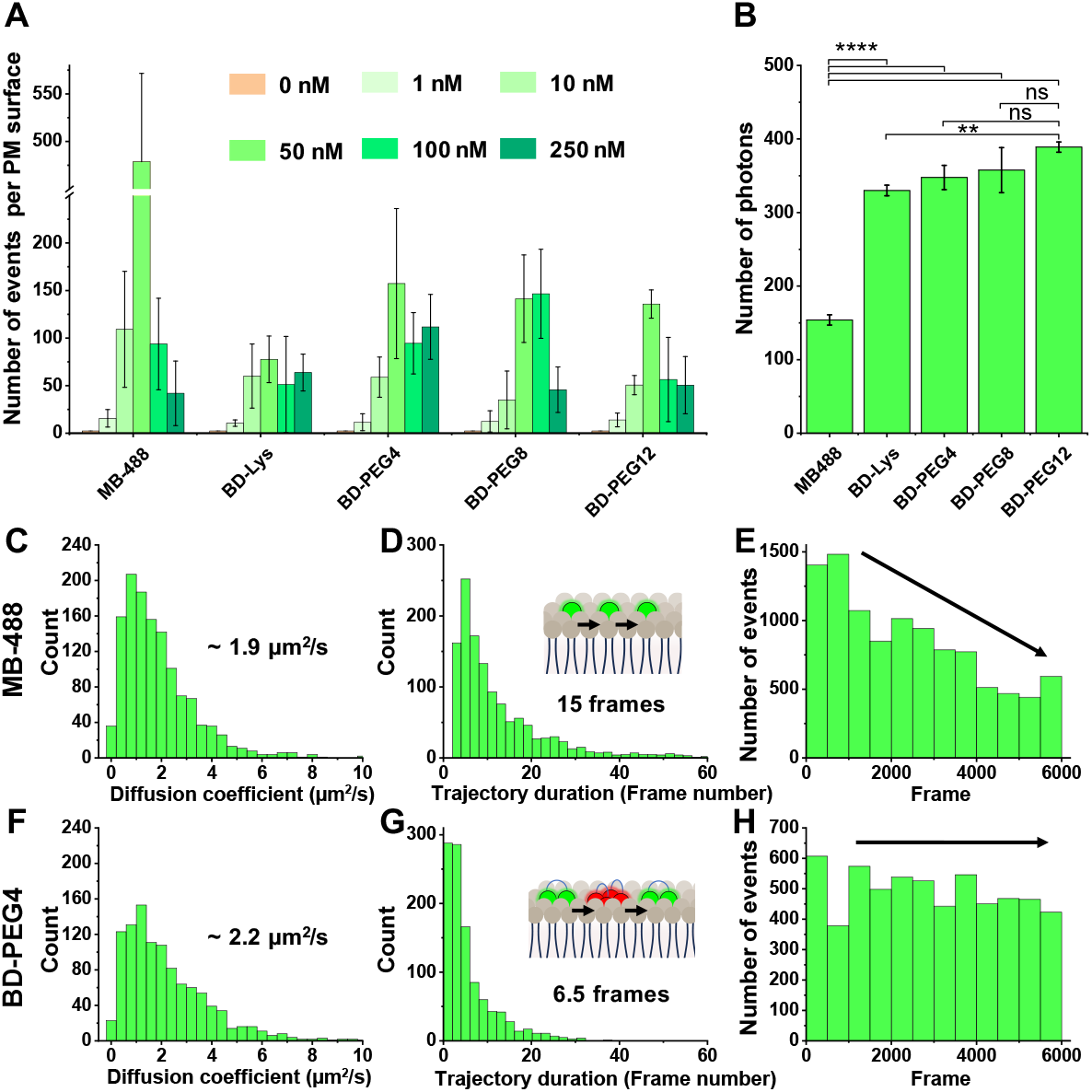
SMLM performance in the green channel. (A) Number of events per PM surface depending on the dye concentrations (0-250 nM) (B) Average number of photons per event for each probe at 50 nM. (C-D and F-G) Histogram of diffusion coefficient and tracking trajectory duration expressed in number of frames for MB-488 and BD-PEG4. (E and H) Histogram of number of events over the acquisition. The measurements were performed on live HeLa cells, excitation laser 488 nm (0.1 kW/cm^2^), 6000 frames, integration time was 13.9 ms. Error bars represent the standard deviation from the analysis of three different cells. ns: not significant. ** : p < 0.01, **** : p < 0.0001, CI: 95%.

To better understand these observations, the monomeric PM probe MB-488 was tested under the same conditions. While 50 nM was also determined as the optimal concentration, the number of events was ≈3-fold higher than that obtained with the dimers.

Intrigued by these results, we first verified that the dimers yielded approximately twice as many photons per event as the monomer (Figure 6B), confirming that each “green” event originated from single fluorophores. Upon careful inspection of the acquisition movies, we noticed that while both types of probes diffused within the membrane and could be tracked over several frames, the dimers exhibited a much more pronounced blinking behavior (See movies in the SI). Quantification confirmed that MB-488 and BD-PEG4 diffuse with similar diffusion coefficients (1.9 vs 2.2 µm^2^/s, respectively), but the monomeric probe could be tracked for ≈15 frames, compared with ≈6.5 frames for the dimer (Figure 6C–D & F–G). These observations confirmed that the dimers are more prone to *J*-aggregation, which intermittently interrupts tracking of the green-emitting open form. Consequently, because MB-488 primarily diffuses in its emissive form, it is more susceptible to photobleaching, and its number of events decreases over 6000 frames, whereas BD-PEG4, which blinks, maintains a nearly constant number of events over time.

Then the obtained reconstructed images were analyzed. At the optimal concentration of 50 nM and after the acquisition of 3500 frames (integration time 13.9 ms), both MB-488 and the dimer BD-PEG4 produced SMLM images with an apparent resolution enhancement of PM segments compared with widefield microscopy (Figure 7A–B). Indeed, when intensity profiles across PM segments were measured, both probes enabled a resolution improvement, reflected by a significant reduction of the FWHM. These results indicate that both probes provide a sufficient number of events to generate high-quality SMLM images in the green channel, but also that the events in this channel mainly arise from diffusion for MB-488, whereas they likely result from a combination of diffusion and blinking for BD-PEG4.

**Figure 7.**
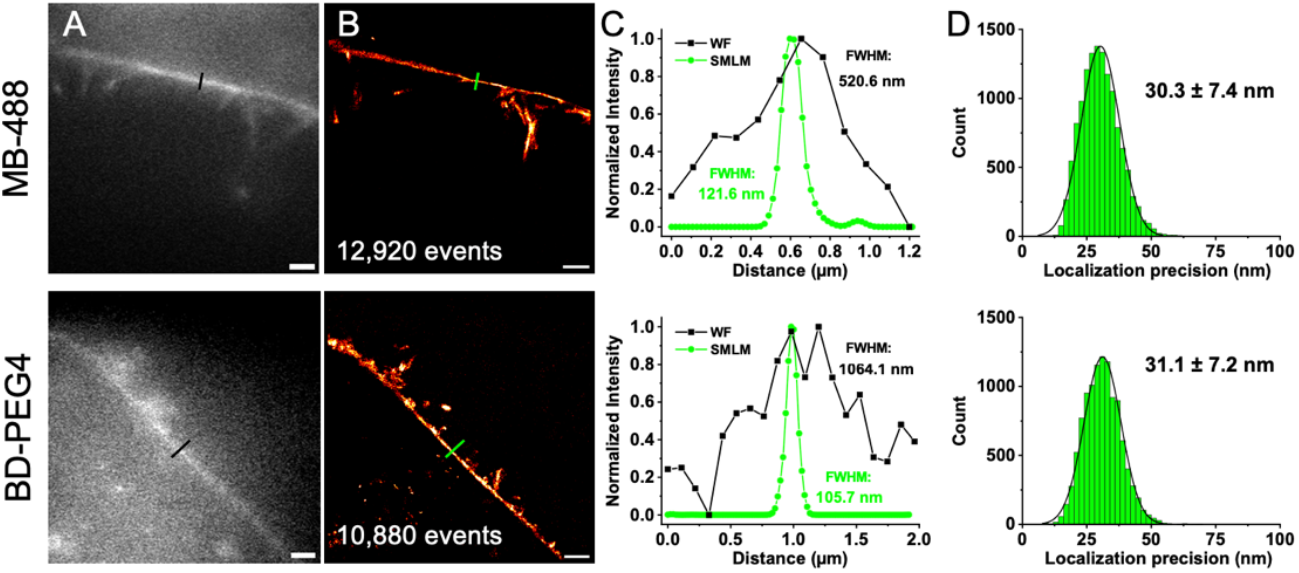
Monomeric MB-488 and dimeric BD-PEG4 provide super resolution images in SMLM in the green channel (λ_ex_= 488 nm). Widefield microscopy images of live HeLa cells’ PM. (B) Reconstructed images from SMLM of the same section than in A obtained from a 3500 frames movie (4:10 min). Probes concentration was 50 nM. Scale bar is 2 µm. (C) Intensity profile corresponding to the black and green line in A and B respectively showing the gain of resolution obtained from SMLM. (D) Localization precision from the localized event. Imaging was performed on live HeLa cells.

Then, to evaluate their performance in the red channel, the probes were incubated at different concentrations with HeLa cells and imaged with an excitation wavelength of 532 nm, at which green-emissive BODIPYs are no longer excited (Figure 3A) and where emissive *J*-aggregates were shown to absorb (Figure S4). While the number of events increased linearly at low probe concentrations (from 0 to 10 nM), it markedly increased and remained rather stable above 50 nM (Figure 8A), which is consistent with the observations of Adhikari et al.^41^ Surprisingly, MB-488 provided the highest number of events per PM surface compared with the dimers, which displayed comparable performances. As observed in the green channel, the number of photons per event was significantly lower for the monomer than for the dimers. These results suggest that *J*-aggregates are more likely to arise from collisions between probes, and thus originate from intermolecular interactions: a collision between two monomers leads to two *J*-aggregated dyes, whereas a collision between two dimers leads to four potential *J*-aggregated fluorophores. This conclusions are consistent with previous studies showing that emissive BODIPY dimers arise primarily from intermolecular interactions within lipid bilayers, and are strongly favored in membrane environments where the local probe density transiently increases.^48^

**Figure 8.**
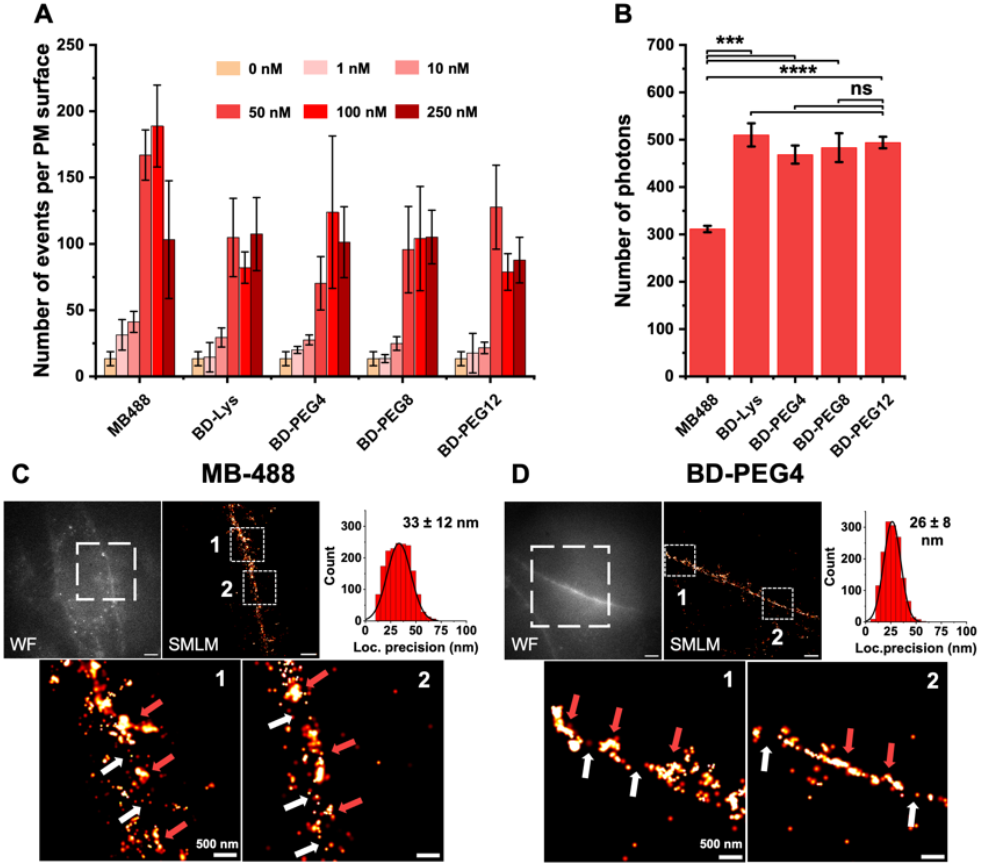
Red-channel SMLM performance of MB-488 and BDs. (A) number of events per PM surface depending on the probes concentrations (0-250 nM). (B) Average number of photons per event for each probe at 50 nM. (C-D) Widefield microscopy and reconstructed images from SMLM of live HeLa cells’ PM section, and the corresponding histogram of localization precision per events. Scale bar is 2 µm. The images in the bottom correspond to the region of interest indicated by the white frame on the widefield microscopy images. Probes concentration was 100 nM. Excitation laser 532 nm (0.5 kW/cm^2^), 6000 frames, integration time was 13.9 ms. Error bars represent the standard deviation from the analysis of three different cells.

Next, the images obtained with 100 nM of monomeric and dimeric probes were analyzed (Figure 8C–D). The widefield images in the red channel unexpectedly showed a rather high signal-to-noise ratio, particularly in the case of the dimer (Figure 8D), suggesting that emissive *J*-aggregates are formed in substantial amounts or in a sufficiently non-transient manner to be directly visualized. The corresponding SMLM images indicated that, while the events were selectively localized on the PM, the signal was less homogeneous than that observed in the green channel (Figure 7B). Although the SMLM images provided comparable results, BD-PEG4 led to a higher localization precision than the monomer MB-488, which is in line with Figure 8B. Upon zooming into regions of interest, different PM zones could be distinguished: regions composed of highly concentrated and intense events (red arrows) and regions with sparse, dim events (white arrows). This difference between the green and red channels combined with the localized nature of the red-channel signal, suggests that *J*-aggregate formation could be favored in specific PM microdomains such as lipid rafts.

To sum up, diffusion of the green-emitting probes within the PM enabled SMLM imaging with an enhanced resolution. In the red channel, both monomeric and dimeric probes blinked through *J*-aggregation in localized domains. Overall, these results suggest that while the lipid bilayer prevents the formation of *H*-aggregates, it provides a favorable environment for the formation of *J*-aggregates. Surprisingly, the dimeric probes did not yield more events in the red channel than the monomeric probe, suggesting that intramolecular aggregation of dimers does not occur in the PM, and that *J*-aggregation is instead promoted by intermolecular interactions.

To verify this hypothesis, we sought to compare a monomeric and a dimeric probe localized at the cell surface but embedded in an environment distinct from the lipid bilayer. To this end, we synthesized two HaloTag-compatible probes: the monomeric BODIPY BM-Halo and the dimeric BODIPY BD-Halo (Figure 9A, for synthesis see scheme S1). We intended to localize these probes at a HaloTag fused to the platelet-derived growth factor receptor (PDGFR-Halo),^48^ so that they would be exposed to the extracellular environment and unable to diffuse within the plasma-membrane lipid bilayer, thereby limiting probe–probe encounters. This configuration was expected to prevent aggregate formation for the monomer, while the dimer would be capable of forming *J*-aggregates exclusively through intramolecular interactions (Figure 9B). HeLa cells were thus transfected with the corresponding plasmid, incubated with the Halo probes, and washed with serum to remove nonspecific binding. After verifying by confocal microscopy that the probes were indeed localized at the cell surface (Figure S8), SMLM imaging was performed for both probes in the green and red channels (Figure 9C and D).

**Figure 9.**
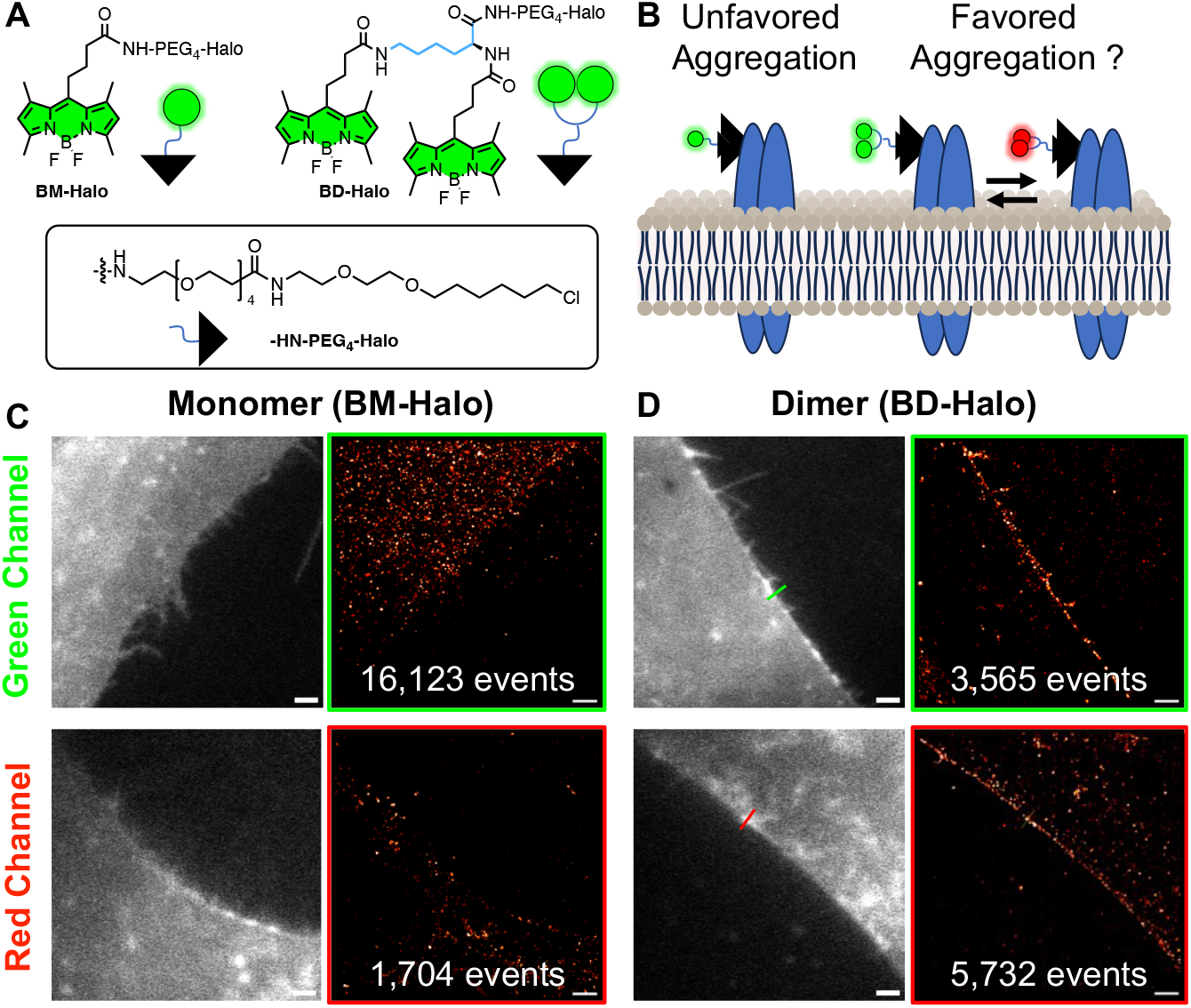
Assessment of the spontaneous blinking properties of monomeric and dimeric HaloTag BODIPYs when localized at the cell surface PDGFR protein and out of the PM bilayer. (A) structure of the designed monomeric and dimeric HaloTag BODIPYs BH-Halo and BD-Halo. (B) Schematic representation of the Halo-probes when bond to the cell surface PDGFR protein depicting the restricted diffusion of probes. (C-D) Widefield and SMLM reconstructed images of transfected HeLa cells labeled with the Halo-probes (50 nM) in both green and red channels. Settings for the green channel: 488 nm (0.2 kW/cm^2^), 8000 frames, integration time was 13.9 ms. and for the red channel: 532 nm (0.5 kW/cm^2^), 8000 frames, integration time was 13.9 ms. Error bars represent the standard deviation from the analysis of three different cells.

Unlike its PM-targeted analogue (MB-488), the monomeric Halo probe BM-Halo only produced off-target events in both green and red channel, in clear discrepancy with the corresponding widefield images, ultimately failing to reconstruct PM sections in either channel (Figure 9C). Although the dimeric Halo probe BD-Halo also generated some off-target events, in some rare cases (Figure 9D) the PM could nonetheless be partially reconstructed in both channels. These results indicate that, at the cell surface but outside the lipid bilayer, BODIPY dimers can still generate detectable events in both green and red channels, likely through transient *J*-aggregation, but to a much lower extent than in the PM.

## Conclusion

In this study, we synthesized plasma membrane–targeted BODIPY dimers, anticipating that spontaneous and transient *H*- and *J*-aggregation would be favored compared to their monomeric analogue MB-488. The two BODIPY units were connected through linkers of increasing length to promote orientations conducive to both types of aggregation. After confirming that the fluorogenic and photophysical properties of the dimeric probes were preserved relative to the monomer (Figures 3 and ref.^4^), we evaluated their aggregation behavior.

Aggregation was first assessed on the precursors by gradually increasing the water content in Water/MeOH mixtures. As expected, whereas the monomer showed no aggregation tendency, the dimers began to aggregate even at low water fractions. Importantly, the aggregation patterns differed markedly among them. The most hydrophobic dimer (p-BD-Lys) first formed intramolecular *H*-aggregates, followed at higher water content by non-emissive intermolecular aggregates (Figure 4). In contrast, the most hydrophilic dimer (p-BD-PEG12) exclusively formed intramolecular *H*-aggregates, while the intermediate dimers, especially p-BD-PEG4, first produced *H*-aggregates and then displayed thin red-shifted bands in both absorption and emission (λ_abs_/λ_em_ = 532/535 nm), consistent with the formation of emissive *J*-aggregates. Concentration-dependent studies confirmed that *J*-aggregation decreased at low probe concentrations, indicating that it is likely driven by intermolecular interactions (Figure S1). In LUVs, the PM probes all began aggregating at probe/lipid ratios above 1/100. The absence of *H*-aggregation, together with a red-shifted broadening of absorption and emission spectra, indicated the formation of *J*-aggregates, particularly for BD-PEG4.

In live SMLM imaging of HeLa plasma membranes in the green channel, the monomeric probe performed best, likely due to its longer diffusion time within the PM, enabling tracking over ~15 frames on average. Although BD-PEG4 diffused with a similar diffusion coefficient (~2 µm^2^/s), consistent with reported BODIPY values in cell membranes,^10^ single molecules could only be tracked for ~6.5 frames on average. As a result, the monomer produced more localization events, although these decreased over time due to photobleaching. In contrast, the dimer yielded fewer events but maintained a nearly constant event rate over 6000 frames owing to its transient blinking behavior. Both probes generated super-resolved images of live PM sections within 3500 frames, with a clear improvement in resolution (Figure 7). These results highlight that non-blinking, photostable, and diffusive monomeric probes are particularly well suited for live SMLM of the plasma membrane.

We next examined *J*-aggregate formation under SMLM conditions in a red channel. In line with Adhikari et al., who showed that monomeric BODIPYs can form transient intermolecular *J*-aggregates in membranes,^41^ the monomer MB-488 generated a high number of red-shifted events. The dimers also produced red events, although in smaller numbers, but with nearly twice as many detected photons per event, consistent with the formation of *J*-aggregates upon intermolecular collisions within the PM. As a result, the dimer BD-PEG4 afforded improved localization precision. In the red channel, localization events appeared sparse yet intense and clustered in specific regions, suggesting that *J*-aggregation may preferentially occur within distinct membrane domains. Although dedicated studies will be required to confirm this hypothesis, this preferential formation of emissive BODIPY dimers is consistent with observations by Gretskaya *et al*. showing that dimer formation is enhanced in membrane regions enriched in complex lipids such as gangliosides, where increased local probe density and molecular packing favor coplanar arrangements.^48^

Overall, our results indicate that *J*-aggregation is strongly favored within lipid bilayers. To further test this, we synthesized monomeric and dimeric HaloTag-BODIPY conjugates targeted to cell-surface PDGFR proteins, thereby restricting probe diffusion and limiting intermolecular interactions. Outside the lipid bilayer, where free diffusion is limited, the monomeric Halo probe produced almost no events at the membrane, whereas the dimer occasionally produced both green and red events, likely through transient intramolecular *J*-aggregation.

In conclusion, this work demonstrates that plasma membrane probes based on green-emissive BODIPYs can be effectively used for live SMLM imaging in the underexploited green channel, thus enabling future multicolor live SMLM applications. Moreover, our findings reveal that the plasma membrane lipid bilayer provides a favorable environment for the formation of emissive BODIPY *J*-aggregates, which may occur preferentially in disordered membrane domains. Although additional studies are needed, simultaneous acquisition in both green and red channels could allow visualization of the entire plasma membrane together with specific subdomains such as lipid rafts, with the high temporal and spatial resolution offered by live SMLM. We hope that this work will stimulate further investigations into the potential of *J*-aggregation for bioimaging, particularly in live multicolor SMLM.

## ASSOCIATED CONTENT

### Supporting Information

Supplementary figures, movies, synthetical protocol and characterization of products as well as NMR and mass spectra can be found in the supplementary information.

## AUTHOR INFORMATION

### Author Contributions

The manuscript was written through contributions of all authors. / All authors have given approval to the final version of the manuscript.

## ACKNOWLEDGMENT

This work was funded by the French National Research Agency (ANR) 5D-SURE ANR-21-CE42-0015. The authors would like to thank Pr. Arnaud Gauthier for providing the plasmid coding for Halo-tagged PDGFR protein.

