## supplementary information for "Membrane-Promoted J-Aggregation of BODIPY Dimers Enables Spontaneous Blinking in Green and Red Channels for Live-Cell Nanoscopy"

### Supplementary figures

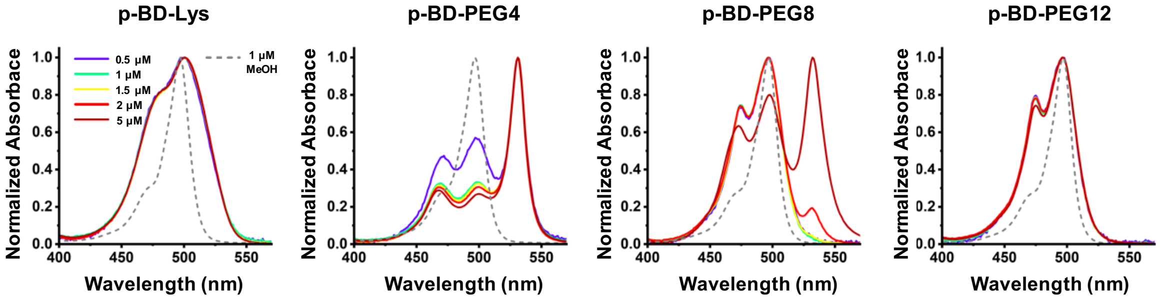


**Figure S1.** Normalized absorption spectra of p-BDs in methanol at 1 µM (dotted line) and in 90/10 water/methanol at increasing concentrations.

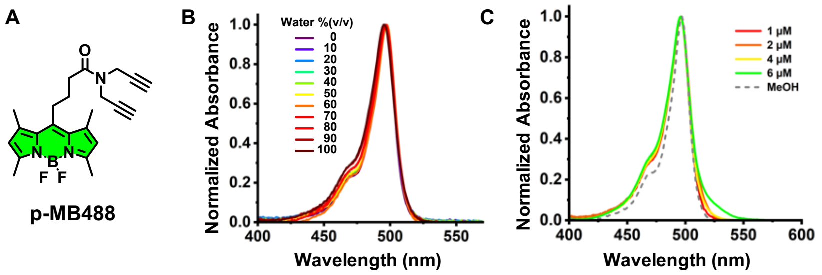


**Figure S2.** (A) Structure of the Membright-488 precursor p-MB488. (B) Normalized absorption spectra of p-MB488 at 1 µM in various Water/MeOH %(v/v). (C) Normalized absorption spectra of p-MB488 in methanol at 1 µM and in 90/10 Water/MeOH %(v/v) upon increasing concentration.

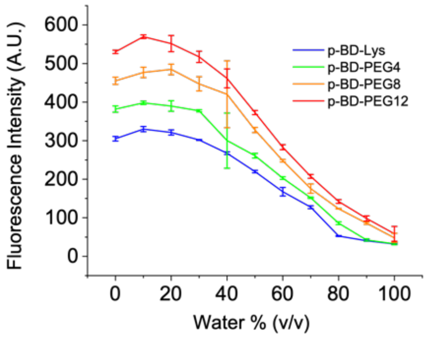


**Figure S3.** Decrease of fluorescence intensity (@ $\lambda_{em\ max}$) of p-BDs (1 µM) upon increase of water/methanol % (v/v).

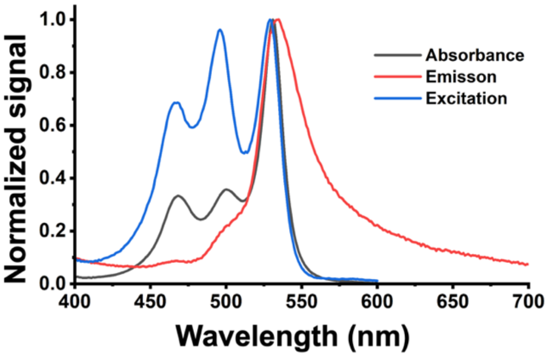


**Figure S4.** Absorbance, excitation and emission spectra of p-BD-PEG4 at 1 µM in 90 % water (v/v). The excitation spectrum was performed at an emission wavelength of 610 nm to minimize the contribution of the remaining bright open form. The emission spectrum was reconstructed form two emission spectra (see methods).

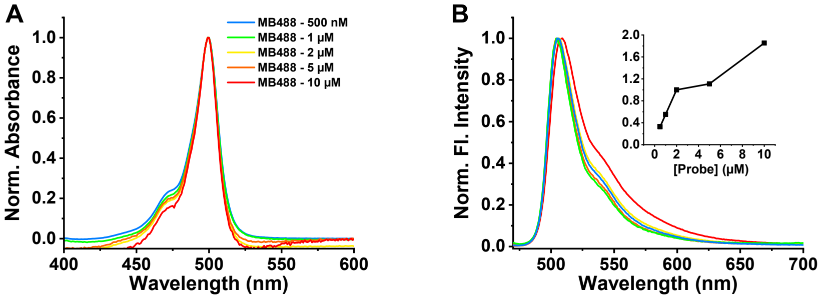


**Figure S5.** Normalized absorption (A) and emission (B) spectra of MB-488 PM probe once inserted in DOPC LUVs (in PBS pH 7.4) at various probe/lipid ratio to increase its local membrane concentration. Inset in B is the normalized fluorescence intensity against increasing concentration of MB-488.

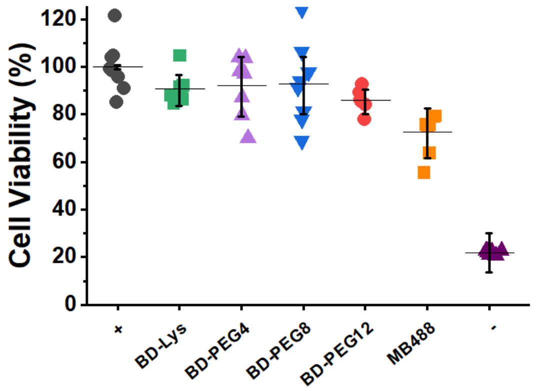


**Figure S6.** MTT cell viability assay for the 4 dimers PM probes and MB-488 (200 nM) after 30 min incubation on HeLa cells. positive control (+) is in absence of probes and with 0.1% v/v DMSO and negative control was obtained by adding 0.1% v/v Triton-X. *: $p < 0.05$. CI: 95%

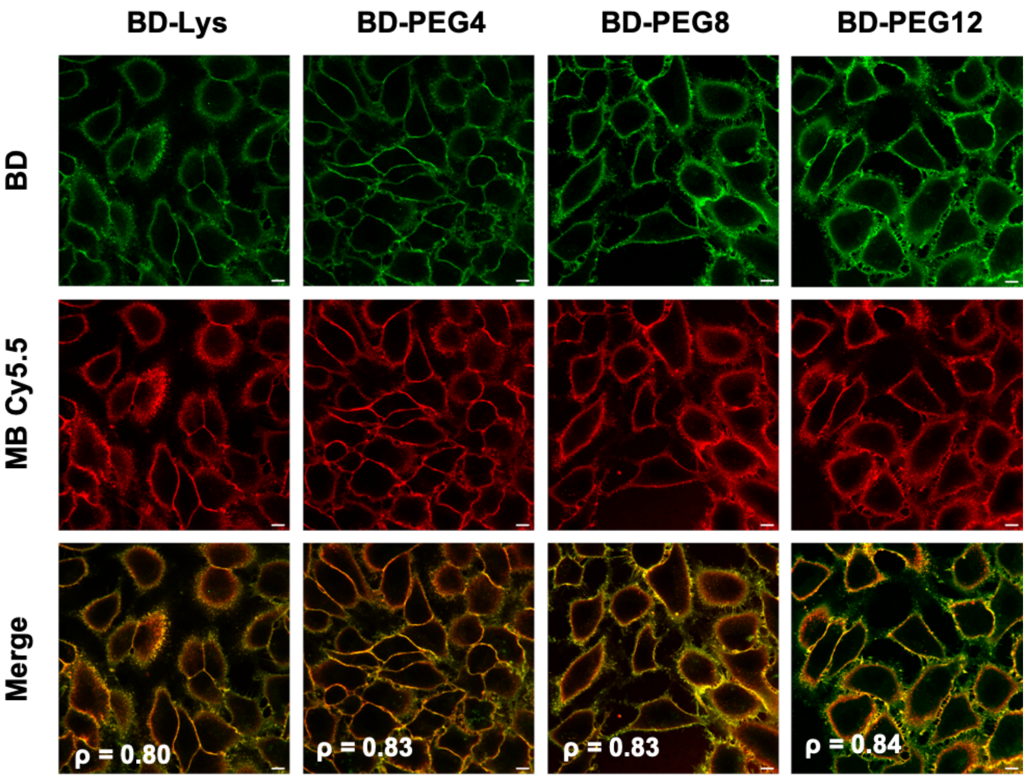


**Figure S7.** Laser scanning confocal microcopy images of HeLa cells 10 minutes after addition of BDs (200 nM) and without washing step. The probes were colocalized with MemBright Cy5.5 confirming their plasma membrane localization. Numbers in white are the Pearson coefficients of the colocalization. Scale bar is 10 µm.

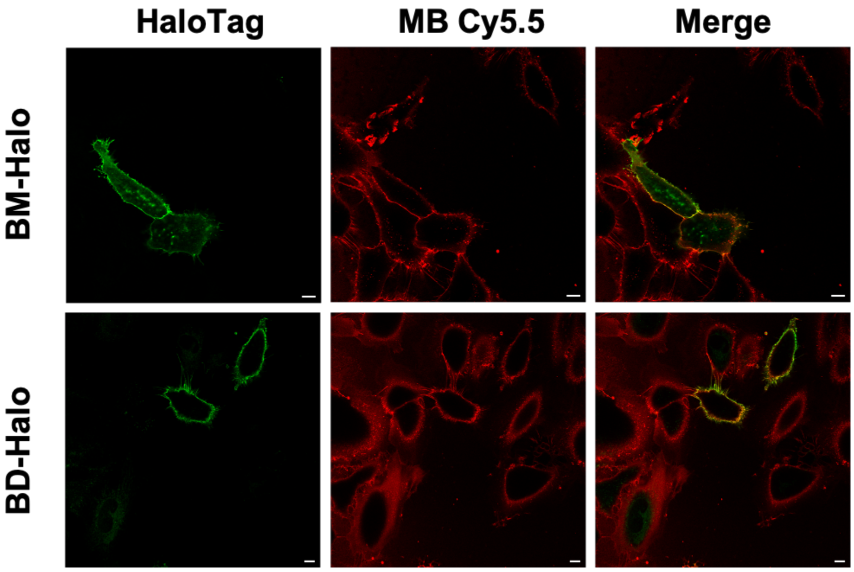


**Figure S8.** Laser scanning confocal microscopy images of live HeLa cells transfected with plasmid coding for Halo-tagged PDGFR and incubated with BM-Halo and BD-Halo (200 nM). MemBright-Cy5.5 (red channel) was used to confirm the localization of the probes at the PM. Scale bar is 10 µm.

### Materials and methods

**Synthesis.** All starting materials for synthesis were purchased from Merck (Darmstadt, Germany), BLD Pharm (Reinbeck, Germany) or TCI Europe (Zwijndrecht, Belgium) and used as received unless stated otherwise. NMR spectra were recorded on a Bruker Avance III 400 MHz or 500 MHz spectrometers. Mass spectra were obtained using an Agilent Q-TOF 6520 mass spectrometer.

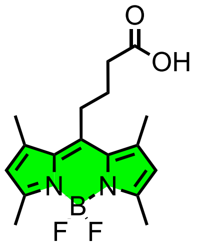


**BDP-COOH**

**BDP-COOH** 4-(4,4-Difluoro-1,3,5,7-tetraMethyl-4-bora-3a,4a-diaza-s-indacen-8-yl)-butyric acid was synthesized according to the literature.[1]

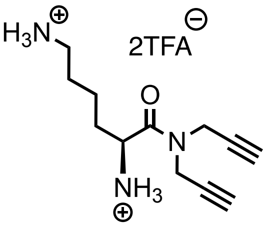


**1**

Di-Boc-L-Lysine dicyclohexylammonium salt (500 mg, 0.95 mmol, 1 eq) was dissolved in DMF (20 mL). HATU (432 mg, 1.14 mmol, 1.2 eq.) was added, followed by dipropargylamine (116 µL, 1.14 mmol, 1.2 eq.) and DIEA (492 µL, 2.84 mmol, 3 eq.). The reaction mixture was stirred at room temperature for 2 hours and then diluted with DCM and water. The aqueous phase was extracted with DCM (3x). The organic phases were combined and dried over $MgSO_4$. The solvent was evaporated in vacuo and the product purified by flash chromatography (50/50 to 70/30 EtOAc/DCM) to give a yellow oil. 2 mL TFA/DCM (1:1) was added and the solution was stirred at room temperature for 1 hour. The solvents were evaporated to give the yellow oil **1** (311 mg, 79%).

$^1$H NMR (400 MHz, MeOD) δ (ppm) 4.55 – 4.22 (m, 5H, $C\underline{H_2}$-N-$C\underline{H_2}$, $\underline{H}$-C-$NH_2$), 2.98 – 2.89 (m, 3H, $C\underline{H_2}$-$NH_3$, C≡C$\underline{H}$), 2.74 (t, $J$ = 2.5 Hz, 1H, C≡C$\underline{H}$), 2.06 – 1.86 (m, 2H, $C\underline{H_2}$-CH-$NH_2$), 1.77 – 1.64 (m, 2H, $NH_2$-$CH_2$-$C\underline{H_2}$), 1.61 – 1.44 (m, 2H, $NH_2$-$CH_2$-$CH_2$-$C\underline{H_2}$).

$^{13}$C NMR (126 MHz, MeOD) δ (ppm) 169.48 ($\underline{C}$=O), 78.30 (C≡C), 78.22 (C≡C), 75.72 (C≡C), 74.33 (C≡C), 51.93 ($\underline{C}$-N), 40.24 ($\underline{C}$-N), 37.24 ($\underline{C}$-$NH_2$), 35.62 ($\underline{C}$-$NH_2$), 31.20 ($C_{aliphatic}$), 28.12 ($C_{aliphatic}$), 22.45 ($C_{aliphatic}$).

HRMS ($ESI^+$): *m/z* calculated for $C_{12}H_{19}N_3O$: 221.1528, found $[M+H]^+$ 222.1607.

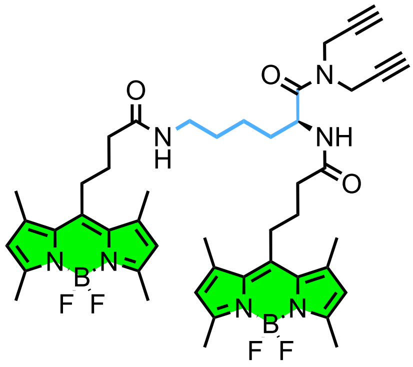

**p-BD-Lys**

To a solution of **BDP-COOH** (50 mg, 0.15 mmol, 1 eq.) and **1** (52 mg, 0.12 mmol, 0.8 eq.) dissolved in DMF (5 mL) was added HATU (68 mg, 0.180 mmol, 1.2 eq.) followed by DIEA (78 µL, 0.448 mmol, 3 eq.). The reaction mixture was stirred at room temperature for 2 hours and then diluted with DCM and water. The aqueous phase was extracted with DCM (3 x). The organic phases were combined and dried over $MgSO_4$. The solvent was evaporated in vacuo and the product was purified by flash chromatography (98/2 to 90/10 DCM/MeOH) to give the orange solid p-BD-Lys (38 mg, 30%).

$^1$H NMR (400 MHz, DMSO-$d_6$) δ (ppm) 8.36 (d, $J$ = 8.2 Hz, 1H, C-N<u>H</u>), 7.90 (t, $J$ = 5.6 Hz, 1H, C-N<u>H</u>), 6.21 (d, $J$ = 2.9 Hz, 4H, C-$\underline{H_{\beta}}$), 4.70 (s, 1H, NH-C<u>H</u>), 4.58 – 4.46 (m, 1H, NH-C<u>H</u>), 4.23 – 4.11 (m, 4H, N-$C\underline{H_2}$), 3.31 (m, 4H, C-$\underline{H_{aliphatic}}$), 3.06 – 2.99 (m, 2H, C≡C<u>H</u>), 2.94 – 2.88 (m, 4H, C-$\underline{H_{aliphatic}}$), 2.39 (s, 24H, C-$\underline{H_{\alpha,\beta}}$), 2.32 (d, $J$ = 7.0 Hz, 2H, C-$\underline{H_{aliphatic}}$), 2.24 (t, $J$ = 6.9 Hz, 2H, C-$\underline{H_{aliphatic}}$), 1.75 (m, 4H, C-$\underline{H_{aliphatic}}$), 1.67 (m, 1H, C-$\underline{H_{aliphatic}}$), 1.52 (m, 1H, C-$\underline{H_{aliphatic}}$), 1.40 (d, $J$ = 7.2 Hz, 2H, C-$\underline{H_{aliphatic}}$).

$^{13}$C NMR (126 MHz, DMSO-$d_6$) δ (ppm) 171.25 (C=O), 170.98 (C=O), 153.11 (C=O), 146.29 ($C_{aromatic}$), 146.17 ($C_{aromatic}$), 140.93 ($C_{aromatic}$), 130.71 ($C_{aromatic}$), 121.66 ($C_{aromatic}$), 78.91 (C≡C), 75.40 (C≡C), 74.59 (C≡C), 48.56 ($C_{aliphatic}$), 36.17 ($C_{aliphatic}$), 35.41 ($C_{aliphatic}$), 34.89 ($C_{aliphatic}$), 34.27 ($C_{aliphatic}$), 31.11 ($C_{aliphatic}$), 28.81 ($C_{aliphatic}$), 27.57 ($C_{aliphatic}$), 27.13 ($C_{aliphatic}$), 27.06 ($C_{aliphatic}$), 22.69 ($C_{aliphatic}$), 15.81 ($C_{aliphatic}$), 15.77 ($C_{aliphatic}$), 14.04 ($C_{aliphatic}$).

$^{11}$B NMR (128 MHz, DMSO-$d_6$) δ (ppm) 0.37 (t, $J$ = 33.6 Hz).

$^{19}$F NMR (376 MHz, DMSO-$d_6$) δ (ppm) -143.88 (m).

HRMS (ESI$^+$): $m/z$ calculated for $C_{46}H_{57}B_2F_4N_7O_3$: 851.4706, found $[M+Na]^+$ 875.4564.

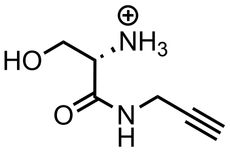

**2**

$N$-Boc-L-Serine (1 g, 4.87 mmol, 1 eq) was dissolved in DMF (30 ml). HATU (222 g, 5.85 mmol, 1.2 eq.) was added, followed by propargylamine (375 µL, 5.85 mmol, 1.2 eq.) and DIEA (2.6 mL, 14.62 mmol, 3 eq.). The reaction mixture was stirred at room temperature overnight, then diluted with DCM and water. The aqueous phase was extracted with DCM (3 x). The organic phases were combined and dried over $MgSO_4$. The solvent was evaporated in vacuo and the product purified by flash chromatography (60/40 to 80/20 EtOAc/Heptane) to give the white solid **2** (1.06 g, 90%). $^1$H NMR is in accordance with the literature.[2]

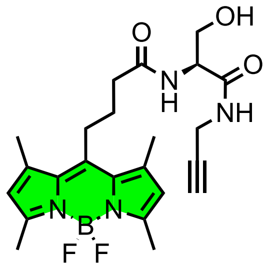

**3**

Product **2** was deprotected by adding 6 mL TFA/DCM (1:1) and stirring for 6 hours. Solvents were evaporated to give a yellow oil **2b** (1.3 g, quant.). 4-(4,4-Difluoro-1,3,5,7-tetraMethyl-4-bora-3a,4a-diaza-s-indacene-8-yl)-butyric acid (300 mg, 0.898 mmol, 1 eq.) was synthesized according to the literature[3] and dissolved in DMF (10 mL). HATU (409 mg, 1.08 mmol, 1.2 eq.) was added, followed by **2b** (279 mg, 1.08 mmol, 1.2 eq.) and DIEA (470 µL, 2.69 mmol, 3 eq.). The reaction mixture was stirred at room temperature for 1 hour, then diluted with DCM and water. The aqueous phase was extracted with DCM (3x). The organic phases were combined and dried over $MgSO_4$. The solvent was evaporated in vacuo and the product purified by flash chromatography (98/2 to 90/10 DCM/MeOH) to give **3** as an orange solid (241 mg, 58%).

$^{1}$H NMR (400 MHz, DMSO-*d*$_6$) δ (ppm) 8.33 (t, *J* = 5.5 Hz, 1H, C-NH), 8.16 (s, 1H ; C-OH), 8.06 (d, *J* = 8.1 Hz, 1H, C-NH), 6.23 (s, 2H, C-$H_{\beta}$), 4.30 (dt, *J* = 8.1, 5.6 Hz, 1H, NH-CH), 3.85 (qdd, *J* = 17.4, 5.8, 2.5 Hz, 2H, NH-$CH_2$), 3.69 – 3.56 (m, 2H, $CH_2$-OH), 3.08 (t, *J* = 2.5 Hz, 1H, C≡CH), 3.03 – 2.91 (m, 2H, C-$H_{aliphatic}$), 2.40 (m, 14H, C-$H_{\alpha,\beta}$ + C-$H_{aliphatic}$), 1.82-1.74 (m, 2H, C-$H_{aliphatic}$).

$^{13}$C NMR (126 MHz, DMSO-*d*$_6$) δ (ppm) 171.46 (C=O), 169.93 (C=O), 153.10 ($C_{aromatic}$), 146.40 ($C_{aromatic}$), 141.04 ($C_{aromatic}$), 130.74 ($C_{aromatic}$), 121.67 ($C_{aromatic}$), 81.05 (C≡C), 72.95 (C≡C), 61.63 (C-O), 54.98 (C-N), 53.60 (C-N), 41.83 ($C_{aliphatic}$), 35.20 ($C_{aliphatic}$), 27.97 ($C_{aliphatic}$), 27.58 ($C_{aliphatic}$), 27.09 ($C_{aliphatic}$).

$^{11}$B NMR (128 MHz, DMSO-*d*$_6$) δ (ppm) 0.38 (t, *J* = 33.0 Hz)

$^{19}$F NMR (376 MHz, DMSO-*d*$_6$) δ (ppm) -69.20, -71.09.

HRMS (ESI$^+$): *m/z* calculated for $C_{23}H_{29}BF_2N_4O_3$: 457.2334, found $[M+Na]^+$ 480.2224.

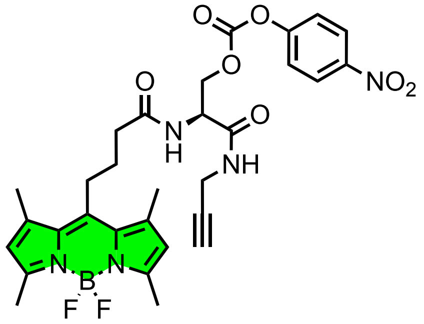

**4**

**3** (200 mg, 0.436 mmol, 1 eq.) was dissolved in anhydrous DCM (10 mL) under argon atmosphere. Pyridine (70 µL, 0.873 mmol, 2 eq.) was added, followed by nitrophenyl chloroformate (145 mg, 0.873 mmol, 2 eq.). The reaction mixture was stirred at room temperature for 40 minutes. The crude mixture was concentrated in vacuo and purified by flash chromatography (98/2 DCM/MeOH) to give **4** as an orange solid (201 mg, 74%).

$^1H$ NMR (400 MHz, DMSO-$d_6$) δ (ppm) 8.67 (t, $J$ = 5.5 Hz, 1H, C-N<u>H</u>), 8.54 (d, $J$ = 8.3 Hz, 1H, C-N<u>H</u>), 8.29-8.24 (m, 2H, $\underline{H}_{ar}$), 7.50-7.45 (m, 2H, $\underline{H}_{ar}$), 6.22 (s, 2H, $C\underline{H}_{\beta}$), 4.77- 4.71 (m, 1H, NH-C<u>H</u>), 4.46-4.36 (m, 2H, NH-$C\underline{H_2}$), 3.97-3.92 (m, 2H, COO-$C\underline{H_2}$), 3.14 (t, $J$ = 2.4 Hz, 1H, C≡C<u>H</u>), 2.95 (dd, $J$ = 7.2, 2.4 Hz, 2H, C-$\underline{H}_{aliphatic}$), 2.43-2.37 (m, 14H, C-$\underline{H}_{\alpha,\beta}$ + C-$\underline{H}_{aliphatic}$), 1.80 (p, $J$ = 6.7 Hz, 2H, C-$\underline{H}_{aliphatic}$).

$^{13}C$ NMR (126 MHz, DMSO-$d_6$) δ (ppm) 181.31 (C=O), 177.51 (C=O), 164.56 (C=O), 162.62 ($C_{aromatic}$), 161.18 ($C_{aromatic}$), 155.61 ($C_{aromatic}$), 154.57 ($C_{aromatic}$), 150.44 ($C_{aromatic}$), 140.20 ($C_{aromatic}$), 134.79 ($C_{aromatic}$), 131.69 ($C_{aromatic}$), 131.13 ($C_{aromatic}$), 90.19 (C≡C), 82.66 (C≡C), 77.51 (C-O), 60.59 (C-N), 44.67 (C-N), 37.65 ($C_{aliphatic}$), 36.83 ($C_{aliphatic}$), 36.52 ($C_{aliphatic}$), 25.27 ($C_{aliphatic}$), 23.51 ($C_{aliphatic}$).

$^{11}B$ NMR (128 MHz, DMSO-$d_6$) δ (ppm) 0.37 (t, $J$ = 33.1 Hz).

$^{19}F$ NMR (376 MHz, DMSO-$d_6$) δ (ppm) -143.91 (p, $J$ = 31.5 Hz).

HRMS ($ESI^+$) : *m/z* calculé pour $C_{30}H_{32}BF_2N_5O_7$: 622.2392, found $[M+Na]^+$ 645.2272.

**General procedure 1.**

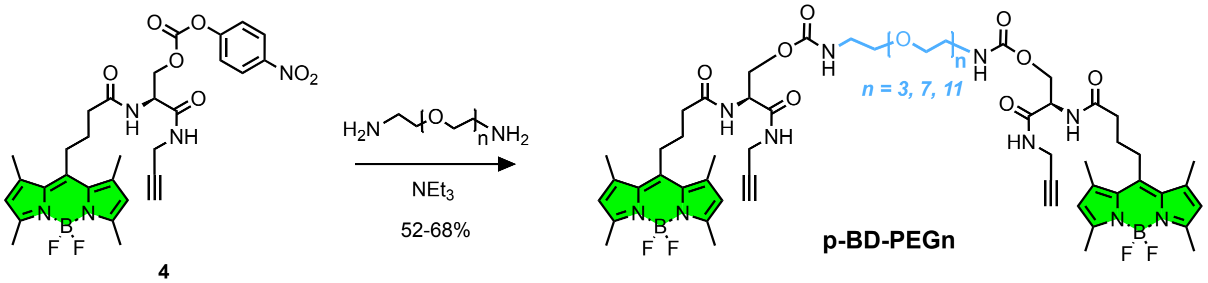


diamino-PEG (1 eq.) was dissolved in DMF (5 mL) and $NEt_3$ was added (2 eq.). Activated carbonate **3** (1.9 eq.) was added to the mixture and the reaction was stirred at room temperature for 3 to 4 hours. The crude mixture was concentrated in vacuo and the products purified by flash chromatography (98/2 to 90/10 DCM/MeOH).

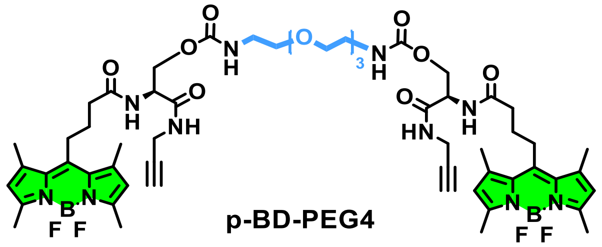


General procedure 1. diamino-PEG : 1,11-diamino-3,6,9-trioxaundecane. **p-BD-PEG4** (26 mg, 60 %).

$^1H$ NMR (400 MHz, MeOD+$CDCl_3$) δ (ppm) 6.06 (s, 4H, $C\underline{H}_{\beta}$), 4.65 (t, $J$ = 5.6 Hz, 2H, NH-C<u>H</u>), 4.27 – 4.21 (m, 4H, NH-$C\underline{H_2}$), 3.96 (t, $J$ = 3.0 Hz, 4H, COO-$C\underline{H_2}$), 3.59 – 3.50 (m, 8H, PEG), 3.45 (t, $J$ = 5.4 Hz, 4H, PEG), 3.23 (t, $J$ = 5.3 Hz, 4H, PEG), 3.04 – 2.93 (m, 4H, C-$\underline{H}_{aliphatic}$), 2.44 (s, 12H, C-$\underline{H}_{\alpha,\beta}$), 2.42 (m, 4H, C-$\underline{H}_{aliphatic}$), 2.40 (s, 12H, C-$\underline{H}_{\alpha,\beta}$), 2.38 (t, $J$ = 2.6 Hz, 2H, C≡C<u>H</u>), 1.97 – 1.86 (m, 4H, C-$\underline{H}_{aliphatic}$).

$^{13}$C NMR (126 MHz, MeOD+$CDCl_3$) δ (ppm) 173.96 (C=O), 170.12 (C=O), 157.46 (C=O), 154.60 ($C_{aromatic}$), 146.29 ($C_{aromatic}$), 141.67 ($C_{aromatic}$), 132.09 ($C_{aromatic}$), 122.37 ($C_{aromatic}$), 79.60 (C≡C), 72.13 (C≡C), 70.96 (C-O), 70.58 (C-O), 70.35 (C-O), 64.63 (C-O), 53.25 (C-N), 41.27 (C-N), 36.37 (C-N), 29.47 ($C_{aliphatic}$), 28.21 ($C_{aliphatic}$), 28.07 ($C_{aliphatic}$), 16.53 ($C_{aliphatic}$), 14.52 ($C_{aliphatic}$).

$^{19}$F NMR (376 MHz, MeOD+$CDCl_3$) δ (ppm) -146.57 (p, *J* = 31.5 Hz).

$^{11}$B NMR (128 MHz, MeOD+$CDCl_3$) δ (ppm) 0.53 (t, *J* = 32.4 Hz).

HRMS (ESI$^+$): *m/z* calculated for $C_{56}H_{74}B_2F_4N_{10}O_{11}$: 1160.5660, found $[M+Na]^+$ 1183.5582.

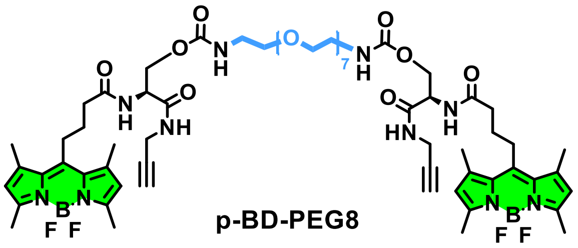

**p-BD-PEG8**

General procedure 1. Diamino-PEG : O,O'-Bis(2-aminoethyl)hexaethylene glycol. **p-BD-PEG8** (19 mg, 52 %).

$^{1}$H NMR (400 MHz, MeOD) δ (ppm) 6.06 (s, 4H, C$\underline{H}_{\beta}$), 4.62 (t, *J* = 5.5 Hz, 2H, NH-C$\underline{H}$), 4.24 (dd, *J* = 5.5, 2.2 Hz, 4H, NH-C$\underline{H_2}$), 3.96 (dd, *J* = 4.9, 2.6 Hz, 4H, COO-C$\underline{H_2}$), 3.60 (d, *J* = 3.0 Hz, 16H, PEG), 3.59 – 3.56 (m, 4H, PEG), 3.55 – 3.52 (m, 4H, PEG), 3.46 (t, *J* = 5.3 Hz, 4H, PEG), 3.23 (t, *J* = 5.7 Hz, 4H, PEG), 3.02 – 2.95 (m, 4H, C-$\underline{H_{aliphatic}}$), 2.46 – 2.39 (m, 28H, C-$\underline{H_{\alpha,\beta}}$ + C-$\underline{H_{aliphatic}}$), 2.37 (d, *J* = 2.7 Hz, 2H, C≡C$\underline{H}$), 1.96 – 1.88 (m, 4H, C-$\underline{H_{aliphatic}}$).

$^{13}$C NMR (126 MHz, MeOD+$CDCl_3$) δ (ppm) 173.80 (C=O), 170.00 (C=O), 157.34 (C=O), 154.51 ($C_{aromatic}$), 146.20 ($C_{aromatic}$), 141.57 ($C_{aromatic}$), 132.02 ($C_{aromatic}$), 122.30 ($C_{aromatic}$), 79.55 (C≡C), 72.05 (C≡C), 70.95 (C-O), 70.93 (C-O), 70.92 (C-O), 70.89 (C-O), 70.59 (C-O), 70.31 (C-O), 64.54 (C-O), 53.17 (C-N), 41.21 (C-N), 36.27 (C-N), 29.42 ($C_{aliphatic}$), 28.07 ($C_{aliphatic}$), 27.98 ($C_{aliphatic}$), 16.51 ($C_{aliphatic}$), 14.52 ($C_{aliphatic}$).

$^{11}$B NMR (128 MHz, MeOD) δ (ppm) 0.53 (t, *J* = 32.9 Hz).

$^{19}$F NMR (376 MHz, MeOD) δ (ppm) -146.47 (p, *J* = 31.5 Hz*)*

HRMS (ESI$^+$): *m/z* calculated for $C_{64}H_{90}B_2F_4N_{10}O_{15}$: 1336.6709, found $[M+Na]^+$ 1359.6576.

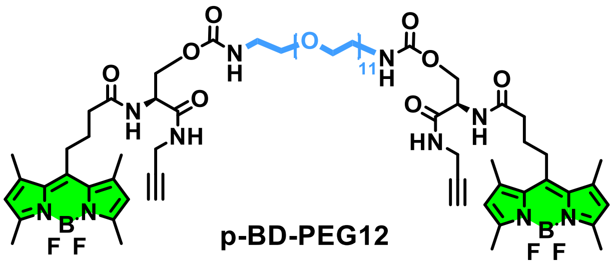

**p-BD-PEG12**

General procedure 1. Diamino-PEG: amino-PEG11-amine. **p-BD-PEG12** (39 mg, 68 %).

$^{1}$H NMR (400 MHz, MeOD) δ (ppm) 6.09 (s, 4H, C$\underline{H}_{\beta}$), 4.64 (t, *J* = 5.6 Hz, 2H, NH-C$\underline{H}$), 4.26 (d, *J* = 5.6 Hz, 4H, NH-C$\underline{H_2}$), 3.96 (dd, *J* = 4.3, 2.6 Hz, 4H, COO-C$\underline{H_2}$), 3.61 (d, *J* = 3.2 Hz, 32H, PEG), 3.59 – 3.56 (m, 4H, PEG), 3.55-3.52 (m, 4H, PEG), 3.45 (t, *J* = 5.4 Hz, 4H, PEG), 3.23 (t, *J* = 5.3 Hz, 4H, PEG), 3.04 – 2.95 (m, 4H, C-$\underline{H_{aliphatic}}$), 2.49 – 2.39 (m, 30H, C-$\underline{H_{\alpha,\beta}}$ + C-$\underline{H_{aliphatic}}$ + C≡C$\underline{H}$), 1.99 – 1.86 (m, 4H, C-$\underline{H_{aliphatic}}$).

$^{13}C$ NMR (126 MHz, MeOD+$CDCl_3$) δ (ppm) 174.31 (C=O), 170.37 (C=O), 157.73 (C=O), 154.75 ($C_{aromatic}$), 146.65 ($C_{aromatic}$), 141.97 ($C_{aromatic}$), 132.30 ($C_{aromatic}$), 122.49 ($C_{aromatic}$), 79.88 (C≡C), 78.94 (C≡C), 78.61 (C-O), 78.29 (C-O), 72.25 (C-O), 71.20 (C-O), 71.14 (C-O), 70.85 (C-O), 70.52 (C-O), 64.71 (C-O), 53.61 (C-N), 41.46 (C-N), 36.47 (C-N), 29.53 ($C_{aliphatic}$), 28.44 ($C_{aliphatic}$), 28.22 ($C_{aliphatic}$), 16.58 ($C_{aliphatic}$), 14.54 ($C_{aliphatic}$).

$^{11}B$ NMR (128 MHz, MeOD) δ (ppm) 0.52 (t, $J$ = 32.7 Hz).

$^{19}F$ NMR (376 MHz, MeOD) δ (ppm) -146.66 (q, $J$ = 31.1 Hz).

HRMS ($ESI^+$): *m/z* calculated for $C_{72}H_{106}B_2F_4N_{10}O_{19}$: 1512.7757, found $[M+H_3O^+]^+$ 1530.8118.

**General procedure 2.**

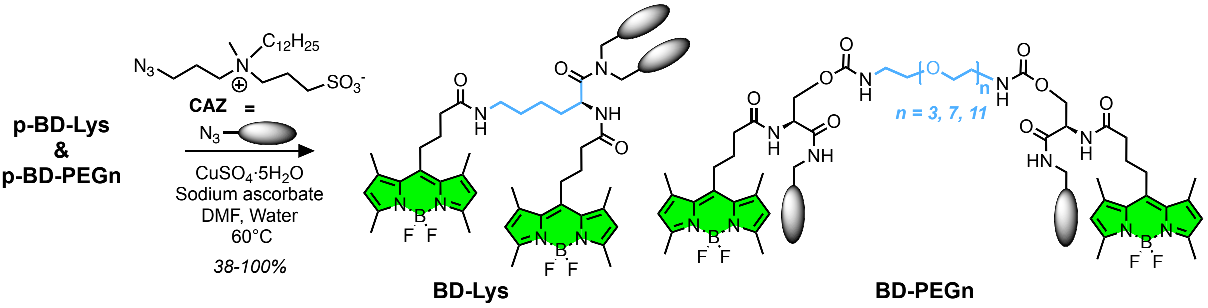


3-((3-azidopropyl)(dodecyl)(methyl)ammonio)propane-1-sulfonate (**CAZ**) was synthesized according to the literature.[4] p-BD-Lys or p-BD-PEGn (1 eq.) was dissolved in DMF (2 mL) and CAZ was added. Copper sulfate pentahydrate (5 mg) and sodium ascorbate (5 mg) were dissolved in an eppendorf with water (200 µL) and the solution vortexed until the mixture turned yellow/orange. The contents of the Eppendorf were then added to the mixture and the reaction stirred at 60°C for 0.5 to 2 hours. The mixture was filtered through celite pad and concentrated in vacuo. The crude product was purified by gel filtration chromatography with DCM/MeOH (1/1) as eluent to obtain an orange solid BD-Lys or BD-PEGn. As the probes are complex and amphiphilic, only their HRMS analyses were carried out.

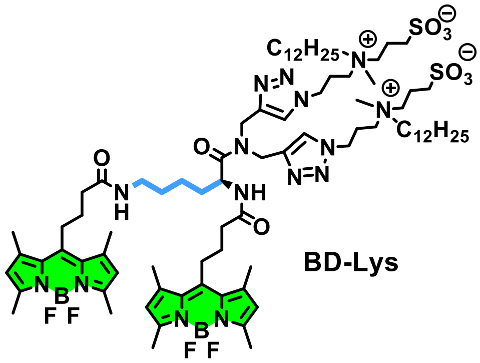

BD-Lys

General procedure 2. BD-Lys (31 mg, 77 %).

HRMS ($ESI^+$): *m/z* calculated for $C_{84}H_{137}B_2F_4N_{15}O_9S_2$: 1662.0287, found $[M+Na]^+$ 1685.0362

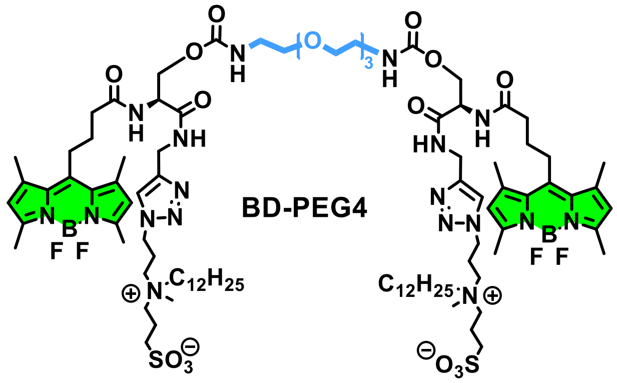


General procedure 2. **BD-PEG4** (10 mg, 38 %).

HRMS (ESI+): *m/z* calculated for $C_{94}H_{154}B_2F_4N_{18}O_{17}S_2$: 1969.1303, found $[M+2H]^{2+}$ 985.5725.

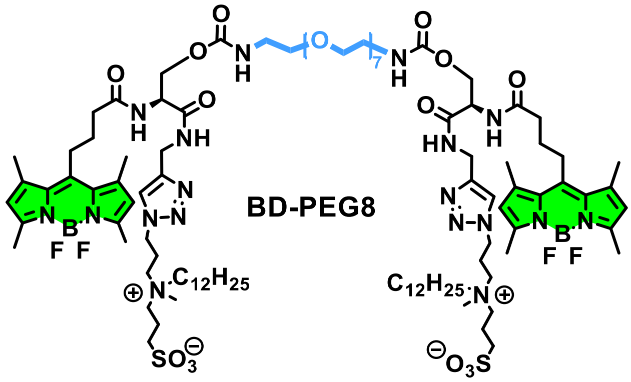


General procedure 2. **BD-PEG8** (25 mg, 103 %).

HRMS (ESI+): *m/z* calculated for $C_{102}H_{170}B_2F_4N_{18}O_{21}S_2$: 2145.2352, found $[M+2H]^{2+}$ 1074.1293.

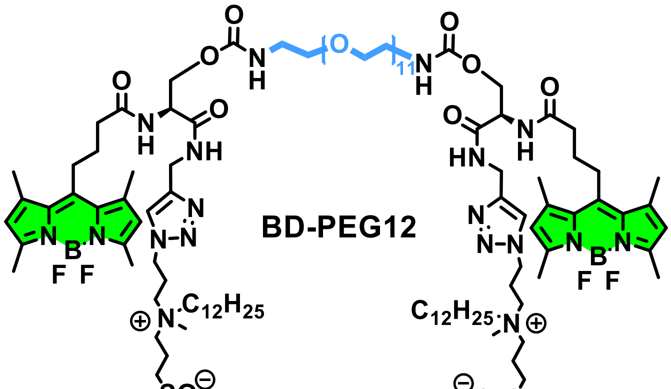


General procedure 2. **BD-PEG12** (15 mg, 49 %).

HRMS (ESI+): *m/z* calculated for $C_{110}H_{186}B_2F_4N_{18}O_{25}S_2$: 2319.3473, found $[M+2Na]^{2+}$ 1183.1605.

**Synthesis of the Halo Probes:**

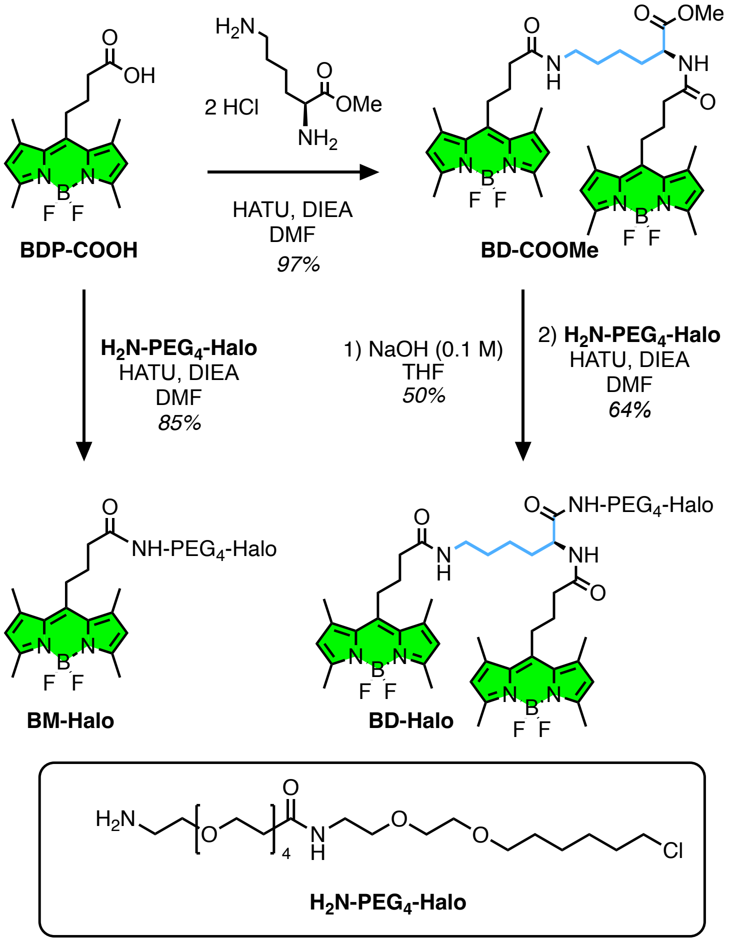


**Scheme S1.** Synthesis of monomeric and dimeric Halo-tagged probes BM-Halo and BD-Halo.

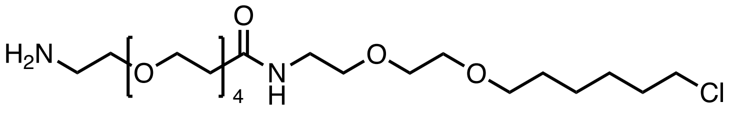

**$H_2N$-PEG$_4$-Halo**

**$H_2N$-PEG$_4$-Halo**, 1-amino-N-(2-(2-((6-chlorohexyl)oxy)ethoxy)ethyl)-3,6,9,12-tetraoxapentadecan-15-amide was synthetized according to the literature.[5]

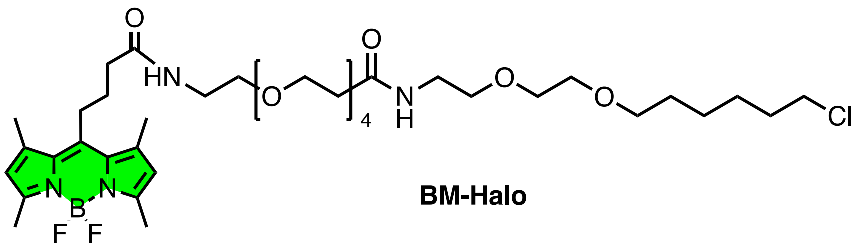

**BM-Halo**

To a solution of BDP-COOH (21.7 mg, 0.07 mmol, 1 eq.) in degassed DMF (5 mL), was added HATU (171 mg, 0.45 mmol, 3 eq.) followed by **$H_2N$-PEG$_4$-Halo** (50 mg, 0.08 mmol, 1.2 eq.) and DIEA (160 µL, 0.9 mmol, 6 eq.). The solvent was evaporated in vacuo and the product was purified by flash chromatography (98/2 to 95/5 DCM/MeOH) to give the orange solid **BM-Halo** (49 mg, 85%).

$^1$H NMR (400 MHz, $CDCl_3$) δ (ppm) 6.69 – 6.40 (m, 2H, 2xN$\underline{H}$), 6.02 (s, 2H, C$\underline{H}_{\beta}$), 3.71 (t, J = 6.0 Hz, 2H, N-C$\underline{H}_2$), 3.63 – 3.48 (m, 22H, PEG), 3.45 – 3.38 (m, 6H, N-C$\underline{H}_2$ + 2 x O-C$\underline{H}_2$), 3.04 – 2.89 (m, 2H, C$\underline{H}_2$-Cl), 2.51 – 2.38 (m, 14H, C-$\underline{H}_{\alpha,\beta}$ + C$\underline{H}_{2,\,meso}$), 2.34 (t, J = 7.0 Hz, 2H, $\underline{H}_{aliphatic}$), 1.92 (q, J = 7.9 Hz, 2H, $\underline{H}_{aliphatic}$), 1.80 – 1.69 (m, 2H, $\underline{H}_{aliphatic}$), 1.57 (p, J = 6.8 Hz, 2H, $\underline{H}_{aliphatic}$), 1.46 – 1.30 (m, 4H, $\underline{H}_{aliphatic}$).

$^{13}$C NMR (126 MHz, $CDCl_3$) δ (ppm) 171.90 (C=O), 171.37 (C=O), 154.05 ($C_{aromatic}$) , 145.67 ($C_{aromatic}$), 140.69 ($C_{aromatic}$), 131.60 ($C_{aromatic}$), 121.79 ($C_{aromatic}$), 71.33 (C-O), 70.58 (C-O), 70.46 (C-O), 70.35 (C-O), 70.30 (C-O), 70.11 (C-O), 69.95 (C-O), 69.87 (C-O), 67.35 (C-O), 45.10 (C-O), 39.42 (C-O), 39.24 (C-O), 37.06 (C-N), 36.27 (C-N), 32.60 (C-Cl), 29.54 ($C_{aliphatic}$), 27.71 ($C_{aliphatic}$), 27.57 ($C_{aliphatic}$), 26.76 ($C_{aliphatic}$), 25.50 ($C_{aliphatic}$), 16.45 ($C_{aliphatic}$), 14.52 ($C_{aliphatic}$).

$^{11}$B NMR (128 MHz, $CDCl_3$) δ (ppm) 0.55 (t, *J* = 32.9 Hz).

$^{19}$F NMR (376 MHz, $CDCl_3$) δ (ppm) -146.46 (p, *J* = 37.9 Hz).

HRMS (ESI$^+$): *m/z* calculated for $C_{38}H_{62}BF_2ClN_4O_8Na$ $[M+Na]^+$: 809.4215, found 809.4229.

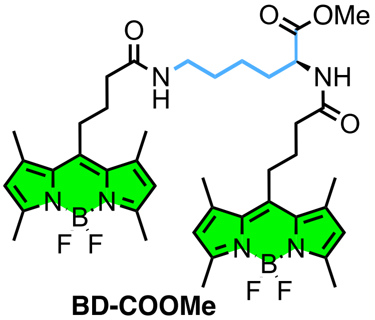


**BD-COOMe**

To a solution of L-lysine methyl ester dihydrochloride (35 mg, 0.15 mmol, 1 eq.) in degassed DMF (5 mL) was added BDP-COOH (125 mg, 0.38 mmol, 2.5 eq.), HATU (171 mg, 0.45 mmol, 3 eq.), and DIEA (160 µL, 0.9 mmol, 6 eq.). The reaction mixture was stirred at room temperature for 2 hours and then diluted with DCM and water. The aqueous phase was extracted with DCM (3 x). The organic phases were combined and dried over $MgSO_4$. The solvent was evaporated in vacuo and the product was purified by flash chromatography (99/1 to 97/3 DCM/MeOH) to give **BD-OMe** as an orange solid (115 mg, 97%).

$^1$H NMR (400 MHz, DMSO-$d_6$) δ (ppm) 8.35 (d, *J* = 7.4 Hz, 1H, N-$\underline{H}$), 7.93 (t, *J* = 5.6 Hz, 1H, N-$\underline{H}$), 6.21 (s, 4H, C$\underline{H}_{\beta}$), 4.30 – 4.11 (m, 1H, CONH-C$\underline{H}$-COOH), 3.60 (s, 3H, O-C$\underline{H}_3$), 3.03 (q, *J* = 6.5 Hz, 2H, NH-C$\underline{H}_2$), 2.96 – 2.88 (m, 4H, C-$\underline{H}_{aliphatic}$), 2.39 (s, 24H, C-$\underline{H}_{\alpha,\beta}$), 2.32 (t, *J* = 7.0 Hz, 2H, C-$\underline{H}_{aliphatic}$), 2.25 (t, *J* = 6.9 Hz, 2H, C-$\underline{H}_{aliphatic}$), 1.81 – 1.72 (m, 4H, C-$\underline{H}_{aliphatic}$), 1.71 – 1.52 (m, 2H, C-$\underline{H}_{aliphatic}$), 1.39 (q, *J* = 7.3 Hz, 2H, C-$\underline{H}_{aliphatic}$), 1.30 (q, *J* = 7.3 Hz, 2H, C-$\underline{H}_{aliphatic}$).

$^{13}$C NMR (126 MHz, DMSO-$d_6$) δ (ppm) 172.91 ($\underline{C}$ONH), 171.76 ($\underline{C}$ONH), 171.19 ($\underline{C}$OOH), 153.29 ($C_{aromatic}$), 146.45 ($C_{aromatic}$), 146.34 ($C_{aromatic}$), 141.08 ($C_{aromatic}$), 130.89 ($C_{aromatic}$), 121.85 ($C_{aromatic}$), 56.17 (C-O), 52.08 (C-N), 51.88 (C-N), 38.39 ($C_{aliphatic}$), 38.36 ($C_{aliphatic}$), 35.59 ($C_{aliphatic}$), 35.05 ($C_{aliphatic}$), 30.71

($C_{aliphatic}$), 28.79 ($C_{aliphatic}$), 27.77 ($C_{aliphatic}$), 27.64 ($C_{aliphatic}$), 27.30 ($C_{aliphatic}$), 27.18 ($C_{aliphatic}$), 23.01 ($C_{aliphatic}$), 18.70 ($C_{aliphatic}$), 15.94 ($C_{aliphatic}$), 14.21 ($C_{aliphatic}$).

$^{11}$B NMR (128 MHz, DMSO-*d*$_6$) δ (ppm) 0.38 (t, *J* = 33.1 Hz).

$^{19}$F NMR (471 MHz, DMSO-*d*$_6$) δ (ppm) -143.93.

HRMS (ESI$^+$): *m/z* calculated for $C_{41}H_{52}B_2F_4N_6O_4$ $[M+Na]^+$: 813.4070, found 813.4279

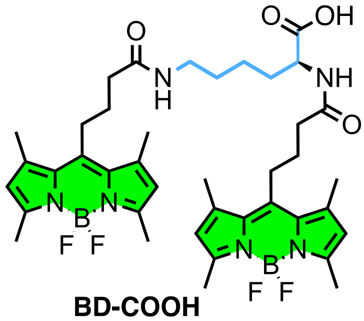


**BD-COOH**

To a solution of **BD-COOMe** (90 mg, 0.11 mmol, 1 eq.) in THF (20 mL) was added aqueous NaOH 0.1 M (1.4 mL, 0.14 mmol, 1.2 eq.) and the reaction mixture was stirred at room temperature for 4 hours. THF was evaporated under vacuum and the crude mixture was diluted with DCM and HCl 1M until pH = 2. The aqueous phase was extracted with DCM (3 x). The organic phases were combined and dried over $MgSO_4$. The solvent was evaporated in vacuo and the product was purified by flash chromatography (95/5 to 85/15 DCM/MeOH) to give **BD-COOH** as an orange solid (44 mg, 50%).

$^1$H NMR (400 MHz, DMSO-d$_6$) δ (ppm) 8.07 (d, J = 7.7 Hz, 1H, N-H), 7.94 (t, J = 5.6 Hz, 1H, N-H), 6.20 (s, 4H, C$H_\beta$), 4.14 – 4.09 (m, 1H, CONH-CH-COOH), 3.02 (q, J = 6.8 Hz, 2H, NH-C$H_2$), 2.97 – 2.85 (m, 4H, C-$H_{aliphatic}$), 2.39 (s, 24H, C-$H_{\alpha,\beta}$), 2.34 – 2.30 (m, 2H, C-$H_{aliphatic}$), 2.24 (t, J = 6.7 Hz, 2H, C-$H_{aliphatic}$), 1.81 – 1.65 (m, 5H, C-$H_{aliphatic}$), 1.56 (q, J = 6.7, 5.6 Hz, 1H, C-$H_{aliphatic}$), 1.42 – 1.35 (m, 2H, C-$H_{aliphatic}$), 1.30 (q, J = 7.3 Hz, 2H, C-$H_{aliphatic}$).

$^{13}$C NMR (126 MHz, DMSO-*d*$_6$) δ (ppm) 171.14 (CONH), 171.00 (CONH), 153.11 (COOH), 146.32 ($C_{aromatic}$), 140.94 ($C_{aromatic}$), 130.73 ($C_{aromatic}$), 121.66 ($C_{aromatic}$), 52.33 (C-N), 48.58 (C-N), 38.39 ($C_{aliphatic}$), 35.41 ($C_{aliphatic}$), 35.18 ($C_{aliphatic}$), 31.13 ($C_{aliphatic}$), 29.00 ($C_{aliphatic}$), 28.79 ($C_{aliphatic}$), 27.65 ($C_{aliphatic}$), 27.59 ($C_{aliphatic}$), 27.14 ($C_{aliphatic}$), 27.09 ($C_{aliphatic}$), 25.47 ($C_{aliphatic}$), 22.95 ($C_{aliphatic}$), 15.81 ($C_{aliphatic}$), 15.77 ($C_{aliphatic}$), 14.04 ($C_{aliphatic}$).

$^{11}$B NMR (128 MHz, DMSO-*d*$_6$) δ (ppm) 0.38 (t, *J* = 33.1 Hz).

$^{19}$F NMR (471 MHz, DMSO-*d*$_6$) δ (ppm) -143.91 (p, *J* = 27.7 Hz).

HRMS (ESI$^+$): *m/z* calculated for $C_{40}H_{51}B_2F_4N_6O_4Na$ $[M+Na]^+$: 800.3991, found 800.4085.

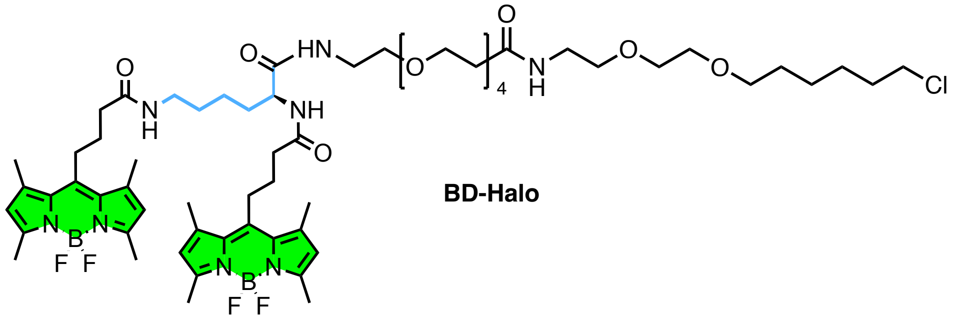


To a solution of **BD-COOH** (15 mg, 0.02 mmol, 1 eq.) in degassed DMF (2 mL) was added: HATU (9 mg, 0.023 mmol, 1.2 eq.), followed by **$H_2N$-$PEG_4$-Halo** (11 mg, 0.023 mmol, 1.2 eq.) and DIEA (10 µL, 0.06 mmol, 3 eq.). The solvent was evaporated in vacuo and the product was purified by flash chromatography (98/2 to 95/5 DCM/MeOH) to give **BD-Halo** as an orange solid (15 mg, 64%).

$^1$H NMR (400 MHz, $CDCl_3$) δ (ppm) 8.51 (s, 1H, N<u>H</u>), 8.23 (d, *J* = 8.3 Hz, 1H, N<u>H</u>), 6.80 (s, 2H, 2xN<u>H</u>), 6.02 (s, 4H, C$\underline{H}_\beta$), 4.41 (q, *J* = 7.3 Hz, 1H, CONH-C<u>H</u>-COOH), 3.73 (t, *J* = 6.0 Hz, 2H, O-C$\underline{H_2}$), 3.59 (d, *J* = 17.3 Hz, 14H, PEG), 3.52 (t, *J* = 6.7 Hz, 6H, PEG), 3.42 (t, *J* = 6.7 Hz, 4H N-C$\underline{H_2}$), 3.20 (p, *J* = 7.4, 7.0 Hz, 2H, N-C$\underline{H_2}$), 3.00 – 2.91 (m, 4H), 2.49 (s, 16H, C-$\underline{H_{\alpha,\beta}}$ + C$\underline{H_{2,\,meso}}$), 2.42 – 2.28 (m, 18H, C-$\underline{H_{\alpha,\beta}}$ + C-$\underline{H_{aliphatic}}$), 1.91 (p, *J* = 6.9 Hz, 4H, $\underline{H_{aliphatic}}$), 1.75 (p, *J* = 6.7 Hz, 3H, $\underline{H_{aliphatic}}$), 1.57 (p, *J* = 6.7 Hz, 3H, $\underline{H_{aliphatic}}$), 1.51 – 1.40 (m, 4H, $\underline{H_{aliphatic}}$), 1.38 – 1.31 (m, 4H, $\underline{H_{aliphatic}}$).

$^{13}$C NMR (126 MHz, $CDCl_3$) δ (ppm) 172.25 (<u>C</u>ONH), 154.23 (<u>C</u>ONH), 145.54 (<u>C</u>ONH), 140.67 (<u>C</u>ONH), 131.60 ($C_{aromatic}$), 128.80 ($C_{aromatic}$), 121.89 ($C_{aromatic}$), 120.29 ($C_{aromatic}$), 71.37 (C-O), 70.59 (C-O), 70.38 (C-O), 70.20 (C-O), 70.11 (C-O), 69.76 (C-O), 67.39 (C-O), 45.16 (C-Cl), 39.38 (C-N), 39.19 (C-N), 36.36 (C-N), 36.32 (C-N), 34.02 ($C_{aliphatic}$), 32.63 ($C_{aliphatic}$), 32.05 ($C_{aliphatic}$), 29.83 ($C_{aliphatic}$), 29.79 ($C_{aliphatic}$), 29.76 ($C_{aliphatic}$), 29.63 ($C_{aliphatic}$), 29.56 ($C_{aliphatic}$), 29.49 ($C_{aliphatic}$), 29.43 ($C_{aliphatic}$), 29.30 ($C_{aliphatic}$), 28.92 ($C_{aliphatic}$), 27.67 ($C_{aliphatic}$), 27.56 ($C_{aliphatic}$), 26.80 ($C_{aliphatic}$), 25.52 ($C_{aliphatic}$), 25.04 ($C_{aliphatic}$), 22.82 ($C_{aliphatic}$), 22.71 ($C_{aliphatic}$), 16.51 ($C_{aliphatic}$), 16.49 ($C_{aliphatic}$), 14.60 ($C_{aliphatic}$), 14.58 ($C_{aliphatic}$), 14.56 ($C_{aliphatic}$), 14.25 ($C_{aliphatic}$).

$^{11}$B NMR (128 MHz, $CDCl_3$) δ (ppm) 0.54 (t, *J* = 33.0 Hz).

$^{19}$F NMR (471 MHz, $CDCl_3$) δ (ppm) -146.48.

HRMS (ESI$^+$): *m/z* calculated for $C_{61}H_{93}B_2ClF_4N_8O_{10}Na$ $[M+Na]^+$: 1253.6723, found 1253.6720.

**Spectroscopy.** Absorption and emission spectra were recorded at 20° C, 1 µM in MeOH on a UV-2700 spectrophotometer (Shimadzu) and a FluoroMax-4 spectrofluorometer (Horiba Jobin Yvon) equipped with a thermo-stated cell compartment. For standard recording of fluorescence spectra, the emission was collected 5 nm after the excitation wavelength. All the spectra were corrected from the wavelength-dependent response of the detector. The fluorescence quantum yields $\phi_F$ was determined by comparison with fluorescein in 0.1M NaOH,[6] as reference according to its excitation and emission wavelengths according to the following equation:

$$\phi_F = \phi_{ref} \times \frac{I_{fluo}^{sample}\, d\lambda}{I_{fluo}^{Ref} d\lambda} \times \frac{OD_{Ref}}{OD_{sample}} \times \frac{n_{sample}^2}{n_{Ref}^2} \quad (1)$$

with $\boldsymbol{I_{fluo}}$: integration of the fluorescence signal, $\boldsymbol{OD}$ : optical density at the excitation wavelength, and $\boldsymbol{n}$ : refraction index of the solvent.

The emission spectrum of the *J*-aggregated BD-PEG4 results from the reconstruction of two emission spectra. The first part of the spectrum was obtained in a "non-conventional" manner by exciting at 560 nm and collecting the fluorescence signal between 400 and 550 nm. The second part was acquired using a conventional approach, with excitation at 525 nm and collection of the emission between 530 and 700 nm. Both emission spectra displayed a maximum intensity at 535 nm; they were therefore normalized and subsequently combined to provide a complete normalized emission spectrum.

**Lipid vesicles.** LUVs were obtained by the extrusion method as previously described.[7] Briefly, a suspension of multilamellar vesicles was extruded by using a Lipex Biomembranes extruder (Vancouver, Canada). The pore size of the filters was first 0.2 µm (10 passages) and thereafter 0.1 µm (10 passages) to generate monodisperse LUVs with a mean diameter of 0.1 µm as measured with a Malvern Zetamaster 300 (Malvern, U.K.). LUVs were composed of DOPC for a final concentration of 200 µM. For the experiment with LUVs, the probe:lipid ratio was set to 1:100, 1:250, 1:500 and 1:1000.

**Cytotoxicity.** Cytotoxicity was evaluated by MTT assay (3-(4,5-dimethylthiazol-2-yl)-2,5-diphenyltetrazolium bromide).[8] A total of $2.10^4$ HeLa cells/well were seeded in a 96-well plate 24 h prior to the experiment in Minimum Essential medium (MEM, Gibco Lifetechnologies) complemented with 10% FBS, Penicillin-Streptomycin (100 µg/mL), L-Glutamine (2 mM), non-essential amino acids (1 mM), sodium pyruvate (1 mM) and were incubated in a 5% $CO_2$ incubator at 37°C. After medium removal, an amount of 100 µL growing media containing 200 nM of Rhod-PM or LRhod-PM were added to the HeLa cells and incubated for 30 min at 37°C (5% $CO_2$). As controls, the cells were incubated with growing media (+ 0.1% DMSO) as positive control and Triton™ X-100 (1%) at negative control of cell viability. After 30 min of dye incubation, the medium was replaced by 100 µL of growing media containing 10 % MTT solution in PBS (4 $mg.mL^{-1}$) and the cells were incubated for 3 h at 37°C (5% $CO_2$). MTT solution was then removed, replaced by 100 µL of DMSO and gently shaken for 10 sec at room temperature. The absorbance at 570 nm was recorded. Each condition was tested in octuplicate, the percentage of cell viability was calculated compared to the control positive control

**Cell sample preparation.** HeLa cells were incubated in Minimum Essential medium (MEM, Gibco Lifetechnologies) complemented with 10% FBS, Penicillin-Streptomycin (100 µg/mL), L-Glutamine (2

mM), non-essential amino acids (1 mM), sodium pyruvate (1 mM) in a 5% $CO_2$ incubator at 37°C. For the imaging experiments, cells were seeded onto a 35 mm IBiDi live cell plates at a density of 90 × $10^3$ cells/well 24 h before the microscopy measurements. 10 minutes prior to acquisition, the culture medium was replaced with 1 mL of Opti-Mem™ containing membrane probes (BD probes, MB Cy5.5).

**Colocalization.** Without any washing steps, 10 minutes after incubation with 200 nM membrane probe (BD, MB or Cy5.5 probes), the cells were imaged using a Leica TSC SP8 confocal laser scanning microscope equipped with a 63x/1.40 objective. The microscope settings were as follows: 488 nm laser (1%) for excitation of BD probes, emission was collected between 500 and 600 nm; 638 nm laser (1%) for MBCy5.5 excitation, with emission collected between 645 and 730 nm. Pearson coefficients were obtained using the ImageJ plugin: Colocalization Finder.

**SMLM imaging.** All samples were imaged using a homemade setup based on a Nikon Eclipse Ti microscope with a 100x 1.49 NA oil immersion objective. An adjustable acousto-optic filter (AOTF; Opto-Electronic) was used to modulate the laser power. The signal was detected with an EM-CCD camera (ImagEM, Hamamatsu, 106 nm per pixel, 5.4 photoelectrons per analog-to-digital conversion unit CAN). For each film, an acquisition time of 13.9 ms per image and 6,000 to 10,000 images were collected.

Green channel: Single molecule images were obtained using 488 nm illumination (max. 0.2 $W.cm^{-2}$) through a dichroic filter (FF497-Di01-25x36, Semrock) and a 488 band-stop filter (NFD01-488-25X36, Semrock). The emission signal is filtered with a bandpass filter (490-540, Olympus).

Red *J*-aggregate channel: Single molecule images were obtained using 532 nm illumination (max. 0.5 $W.cm^{-2}$) through a dichroic filter (Di02-532/635-t1-25x36, Semrock) and a 532 band-stop filter (NF03-532E-25, Semrock). The emission signal was filtered with a bandpass filter (FF01-607/70-25, Semrock).

The SMLM films were analyzed using the ThunderSTORM plug-in of the ImageJ software (version 1.54i). The images were filtered with "Wavelet filter" (B-spline order 3, scale 2.0), the approximate location of the molecules was determined using the local maximum method (intensity threshold of 1.5 × standard deviations, 8-neighbor connectivity), and the sub-pixel location of the molecules was determined using the "maximum likelihood fitting method (PSF integrated Gaussian method, fitting radius 3 px, initial sigma 1.6 px). Data processing was performed by filtering the locations: $20 < \sigma < 200$ and 50 < number of photons < 4000. The reconstructed images were produced using a normalized Gaussian method with a lateral uncertainty of 20 nm. The intensity profiles were adjusted using a Gaussian fit to determine the FWMH values.

**Single particle tracking.** The images used for SPT analyses were produced on samples acquired using the method described for SMLM imaging, with a probe concentration of 50 nM. The films were analyzed using the TrackMate plug-in of the ImageJ software (version 7.12.2.). Molecules were detected using LoG (estimated diameter: 0.6 µm, quality threshold: 250). The trajectory of the molecules was tracked using LAP Tracker (maximum distance: 0.5 µm, maximum image interval between two positions: 2).
